# The Holliday junction resolvase GEN1 preserves genome integrity and self-renewal in mouse embryonic stem cells

**DOI:** 10.64898/2026.07.23.740277

**Authors:** Lucía Ramos-Lage, Cristina Ameneiro, David Martínez-Delgado, Helena Covelo-Molares, Tiago Moreira, Raquel Carreira, Diana Rubio-Contreras, Alba Coego, Vera Garcia-Outeiral, Alejandro Fuentes-Iglesias, Evi Soutoglou, Miguel Fidalgo, Miguel G. Blanco, Diana Guallar

## Abstract

The maintenance of pluripotent stem cells (PSCs) under rapid proliferation requires mechanisms that both suppress replication-driven genome instability and preserve self-renewal capacity. Here, we show that, in contrast to somatic cells where it mainly acts as a backup, the Holliday junction resolvase GEN1 is required in mouse embryonic stem cells (ESCs), where its depletion severely compromises self-renewal and long-term maintenance. Loss of GEN1 induces the accumulation of cells with DNA content greater than 4C and chromosome fusions. Notably, a catalytically inactive GEN1 mutant rescues ESC colony formation, indicating that GEN1 supports ESC maintenance through non-enzymatic functions. In addition, GEN1 depletion increases ESC tolerance to topoisomerase I-mediated replication stress and renders this phenotype dependent on DNA-PK activity, suggesting that GEN1 loss alters how pluripotent cells cope with replication-associated DNA lesions. Together, these findings identify GEN1 as a non-redundant guardian of genome integrity in pluripotent cells, revealing both a catalysis-independent role in self-renewal and a contribution to the replication stress response, with implications for PSC genomic quality control.

**Highlights:**

- In contrast to somatic cells, GEN1 is specifically required for mouse pluripotent cell self-renewal and expansion *in vitro*.
- GEN1 loss induces accumulation of DNA content greater than 4C and chromosome fusions without loss of core pluripotency markers expression.
- Catalytically inactive GEN1 mutant rescues ESC colony-forming capacity.
- GEN1 depletion increases ESC tolerance to topoisomerase I-mediated replication stress in a DNA-PK-dependent manner

**eTOC:** Ramos-Lage et al. demonstrate that the resolvase GEN1 is essential for mouse embryonic stem cell self-renewal and genome stability. Strikingly, a catalytically dead mutant rescues colony formation, revealing an unexpected non-enzymatic role for GEN1 in pluripotency maintenance.

## Introduction

Pluripotent stem cells (PSCs), including embryonic stem cells (ESCs), combine unlimited self-renewal with the capacity to generate derivatives of all three embryonic germ layers. Maintaining genome integrity is therefore essential both for normal development and for the experimental and translational use of PSCs. A hallmark of ESCs is their rapid proliferation, driven by a shortened cell cycle with an abbreviated G1 phase and constitutive cyclin-dependent kinase activity that bypasses the canonical G1/S checkpoint (Savatier *et al*., 2002; Stead *et al*., 2002; White *et al*., 2005). Although S-phase duration is comparable to somatic cells (White *et al*., 2005; Fujii-Yamamoto *et al*., 2005), their accelerated cycling imposes persistent replication stress that can give rise to DNA damage (Ahuja *et al*., 2016; Banáth *et al*., 2009). To safeguard genome integrity, PSCs rely strongly on high-fidelity homologous recombination (HR) repair during S and G2 phases (Tichy *et al*., 2010), minimizing mutations that could compromise pluripotency or be propagated to daughter cells in the embryo. Consistent with this, core HR factors, including the MRN complex (Luo *et al*., 1999; Zhu *et al*., 2001; Tang *et al*., 2024; Xiao and Weaver, 1997), BRCA1 and BRCA2 (Gowen *et al*., 1996; Hakem *et al*., 1996; Hakem, de la Pompa and Mak, 1998), PALB2 (Rantakari *et al*., 2010), and RAD51 (Tsuzuki *et al*., 1996; Yoon *et al*., 2014), are essential for ESC viability and embryonic development.

While early steps of HR have been studied extensively, much less is known about how late recombination intermediates, particularly Holliday junction (HJ) processing, are processed in pluripotent cells. HJs are secondary DNA structures which are processed by dissolution via the BLM-TOP3A-RMI1/2 complex (Wu and Hickson, 2003) or cleavage by structure-selective endonucleases, such as MUS81-EME1/2 and GEN1 (Wyatt and West, 2014; Lilley, 2017; Dehé and Gaillard, 2017). Deficiencies in BLM (Huang *et al*., 2012) and MUS81 (Hanada *et al*., 2006) impair genome stability in ESCs, but the specific contribution of GEN1 in this context has remained unexplored. In somatic cells, GEN1 operates primarily as a backup resolvase, being largely dispensable under normal conditions, but becomes critical upon MUS81 or SLX1/4 loss and the combined depletion with either nuclease is synthetic lethal (Wang *et al*., 2016; Wechsler, Newman and West, 2011). Whether this hierarchy also applies to pluripotent cells, which replicate rapidly within a distinctive chromatin and cell-cycle environment, has not been established.

Here, we investigate how GEN1 contributes to genome stability and self-renewal in mouse ESCs. Using CRISPR/Cas9-mediated knock-in tagging, loss-of-function and rescue approaches alongside shRNA silencing, we show that GEN1 is essential for colony formation, with depletion inducing accumulation of cells with DNA content greater than 4C, chromosome fusions, and replication stress-specific phenotypes. Strikingly, catalytically inactive GEN1 retains the ability to rescue the colony formation defect, revealing non-redundant, largely non-enzymatic functions for GEN1 in supporting ESCs in culture. Furthermore, the absence of GEN1 impacts the sensitivity of ESCs to topoisomerase I-induced replication stress, suggesting that GEN1 is required for proper handling of DNA lesions generated during replication. These findings identify GEN1 as a genome guardian with a previously unrecognized, essential role in pluripotent stem cell maintenance, revealing a cell state–specific dependency not observed in somatic cells.

## Results

### GEN1 is enriched in ESCs and required for self-renewal

Given the prominent role of homologous recombination in preserving pluripotent stem cell genomes, we first examined how the key Holliday junction (HJ) processing factors BLM, MUS81, and GEN1 are expressed in pluripotent stem cells compared to differentiated cells. Analysis of their expression using data from published RNA-seq (Kaemena *et al*., 2023) showed higher mRNA levels for *Blm* and *Gen1* in mouse ESCs than in mouse embryonic fibroblasts (MEFs) (**Figure S1A**). This observation was confirmed by RT-qPCR (**Figure S1B**). In ESCs, *Blm* is expressed at higher levels than *Gen1* and *Mus81*, with *Gen1* showing higher expression than *Mus81*. Consistent with previous reports in somatic cells favouring dissolution over resolution to limit crossovers (Wechsler, Newman and West, 2011; Garner *et al*., 2013), this pattern suggests that BLM-mediated dissolution predominates.

To test whether these factors are required for ESC self-renewal, we performed CRISPR/Cas9-mediated loss-of-function assays using independent pairs of sgRNAs targeting *Blm*, *Mus81* or *Gen1* (**Figures 1A and 1B**). Colony formation assays (CFA) with alkaline phosphatase (AP) staining revealed that targeting *Gen1* markedly reduced AP-positive (AP^+^) colonies, whereas sgRNAs against *Blm* or *Mus81* had only minor effects under these conditions (**Figures 1C and S1C**). Genotyping of surviving clones in sg*Gen1*-treated cells showed retention of the wild-type *Gen1* allele (**Figures 1D and 1E**), consistent with the idea that these surviving colonies derive from rare ESCs in which *Gen1* was not effectively disrupted.

**Figure 1.**
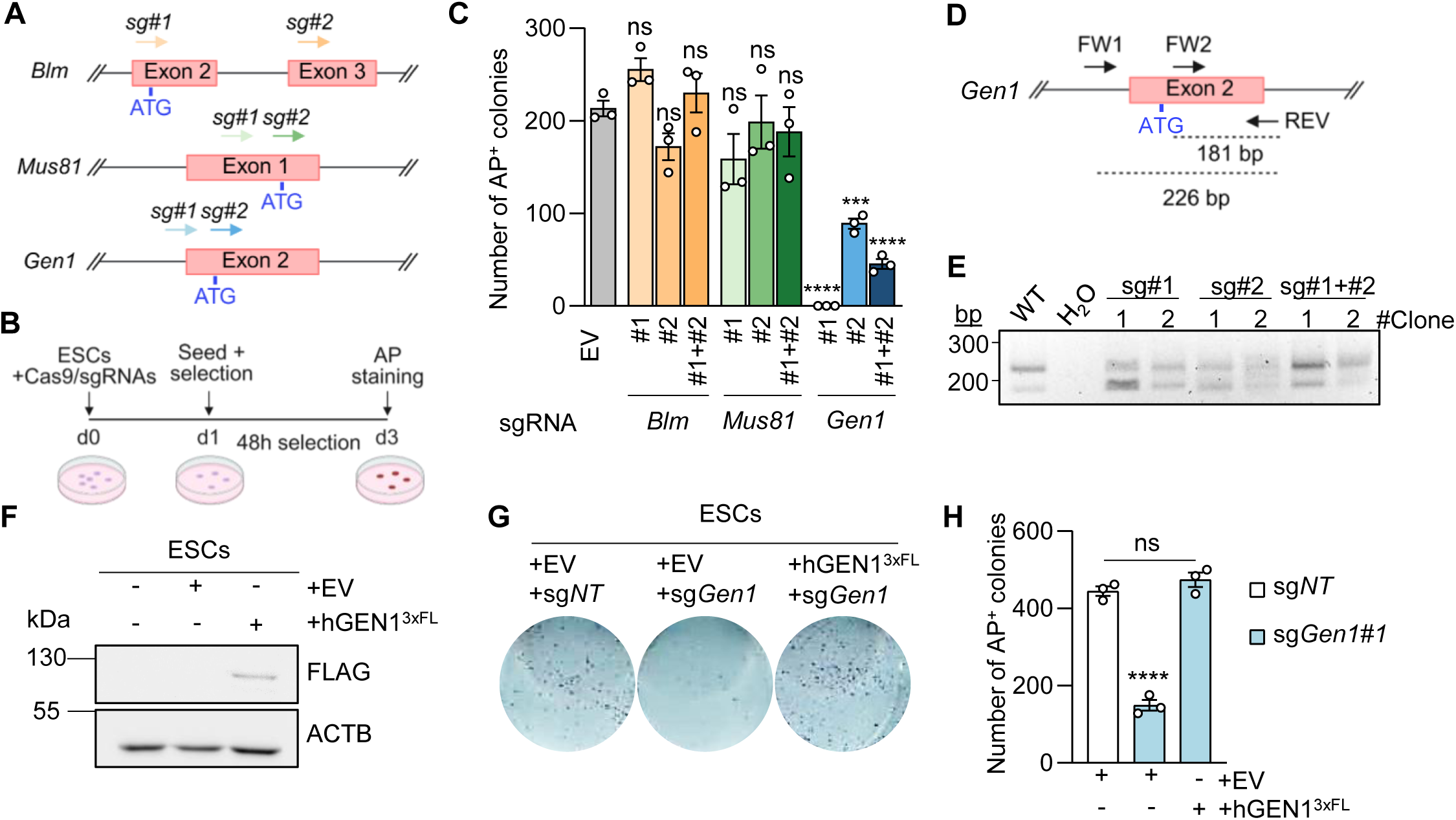
GEN1 is essential for ESC self-renewal. (A) Schematic depiction of the *Blm, Mus81* and *Gen1* genomic *loci* showing the sgRNA target sites (coloured arrows). The translation initiation site (ATG) is indicated for each gene. (B) Strategy for CRISPR/Cas9-mediated knock-out (KO) of *Blm*, *Mus81* and *Gen1* using Cas9-sgRNA plasmids. Cells transfected with sgRNAs targeting the desired genes were seeded for CFA and scored by alkaline phosphatase (AP) staining. (C) Number of AP-positive (AP^+^) colonies obtained under the indicated conditions. Data represent mean ± SEM. (D) Genotyping strategy for verification of *Gen1* KO in ESCs, using a three-primer PCR approach that generates products of 181 bp and 226 bp in the wild type (WT) phenotype. (E) Representative gel showing PCR genotyping products from clones obtained following transfection with sgRNA#1, sgRNA#2, or sgRNA#1 + sgRNA#2 targeting *Gen1*. (F) Western blot showing expression of exogenous human GEN1^3xFL^ detected with an anti-FLAG antibody. ACTB was used as loading control. (G) Representative images of AP staining for each condition indicated. (H) Quantification of AP^+^ colonies formed following transfection with sgRNA targeting *Gen1* or a non-targeting (NT) control, in hGEN1^3×FL^ or EV ESC lines. Data represent mean ± SEM. Statistical significance was assessed by one-way ANOVA with Dunnett’s comparison test to EV conditions. \*\*\**p* < 0.001; \*\*\*\**p* < 0.0001; ns, non-significant.

To confirm that the colony-formation defect was specifically due to *Gen1* loss rather than to off-target effects, we carried out rescue experiments using the human GEN1 ortholog. An ESC line stably overexpressing full-length human GEN1 tagged with 3 FLAG tags (hGEN1^3xFL^) restored AP^+^ colony numbers upon endogenous *Gen1* disruption to levels comparable to non-targeting controls, whereas empty-vector (EV) ESCs showed a strong reduction in AP^+^ colonies in the same setting (**Figures 1F–H**). Together, these findings support a specific requirement for GEN1 in ESC colony formation and suggest that GEN1 functions as an important factor for self-renewal in mouse ESCs.

### GEN1 depletion is associated with chromosome fusions in ESCs

Since stable *Gen1* KO ESC lines could not be maintained, we relied on shRNA-mediated knockdown together with a CRISPR/Cas9 knock-in (KI) ESC line expressing C-terminally 3×FLAG-tagged GEN1 (*Gen1*^3xFL^) to study endogenous GEN1 protein (**Figure S1D-G**). The cell line was generated as a tool to study GEN1 since there is no good commercial antibody against GEN1. Subcellular fractionation and immunofluorescence showed that GEN1 is predominantly cytoplasmic in ESCs, consistent with its reported behaviour in somatic cells (Chan and West, 2014), while a detectable fraction is present in nuclear and chromatin-bound compartments, including after G1/S synchronisation (**Figures S1H and I**), indicating that ESCs retain a nuclear GEN1 pool in interphase.

Colony formation assays confirmed a marked reduction in AP^+^ colonies in sh*Gen1* ESCs compared with sh*Luci* controls (**Figures 2A-C**). We next asked whether *Gen1* loss induces transcriptional alterations underlying this self-renewal defect. The RNA-seq analysis of sh*Gen1* versus sh*Luci* ESCs revealed discrete changes on the transcriptome of the cells, with core pluripotency factor expression unchanged (**Figure S2A**), confirmed as well by RT-qPCR for *Oct4* (**Figure 2D**). In line with this, a gene signature for stem cell population maintenance remained unchanged upon silencing of *Gen1* (**Figure 2E**) These results thus indicate that the self-renewal defect observed upon *Gen1* depletion reflect impaired proliferation rather than identity loss.

**Figure 2.**
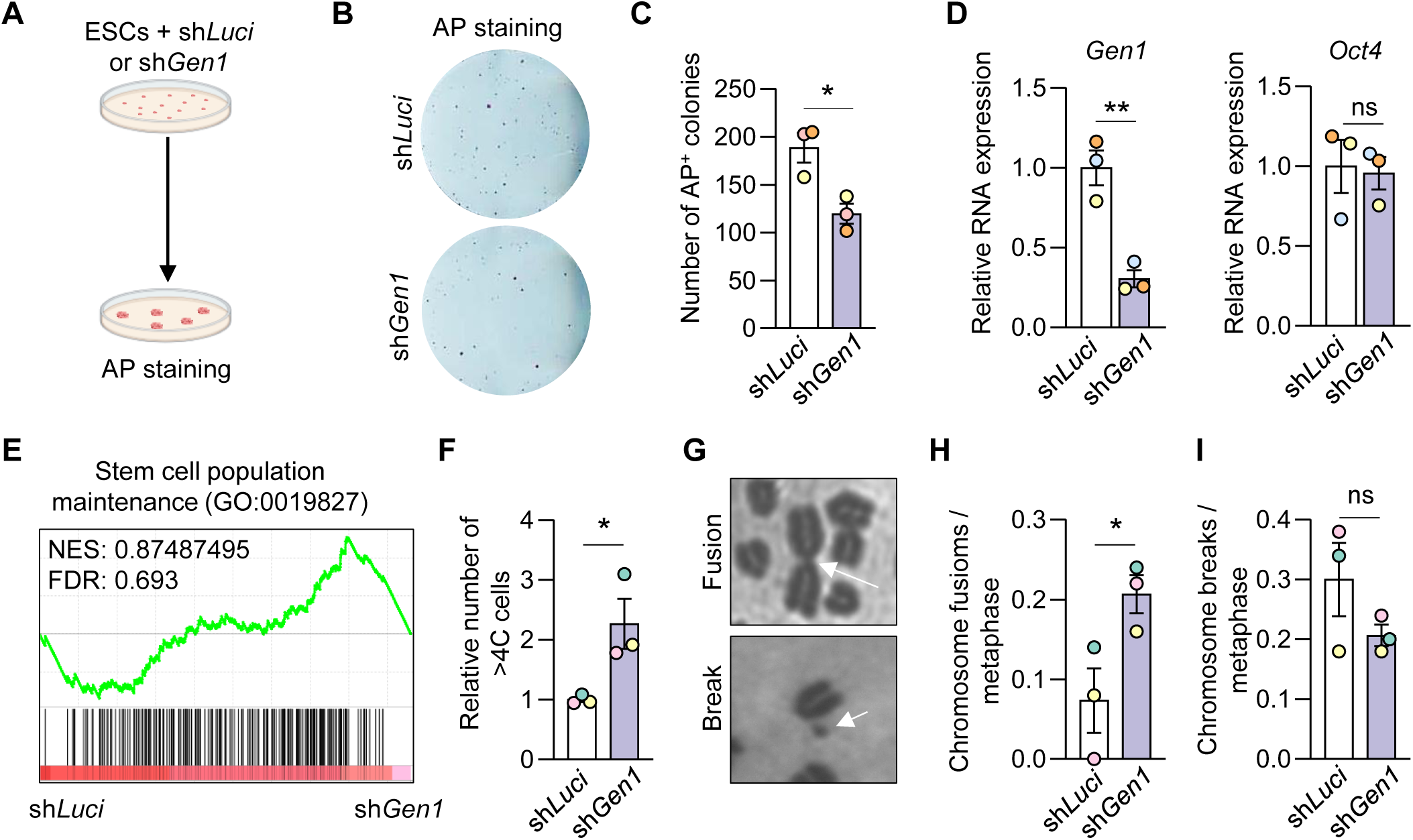
*Gen1* silencing impairs ESC self-renewal capacity and genome stability. (A) Schematic of the experimental approach for *Gen1* silencing in ESCs using shRNAs, with sh*Luci* control. (B) Representative images of wells after AP-staining and (C) quantification of AP^+^ colonies. Data represent mean ± SEM (n = 3, biological replicates). (D) Relative RNA expression of *Gen1* and the pluripotency marker *Oct4* in sh*Gen1* ESCs compared to sh*Luci* as control (n=3, biological replicates). (E) Gene set enrichment analysis from sh*Luci* and sh*Gen1* RNA-seq data against a stem cell population maintenance gene set (GO:0019827). Normalized enrichment score (NES) and FDR are shown. (F) Number of cells with DNA content greater than 4C relative to sh*Luci* (n = 3, biological replicates). Data represent mean ± SEM. (G) Representative images of chromosomal aberrations observed in metaphase spreads: chromosome fusions (upper panel) and chromosome breaks (lower panel), indicated by white arrows. Quantification of chromosome (H) fusions and (I) breaks. Data represent mean ± SEM. Fifty metaphases per condition were scored (n = 3, biological replicates). Statistical significance was assessed by unpaired two-tailed t-test. \**p* < 0.05; \*\**p* < 0.01; ns, non-significant.

On the other hand, cell-cycle analysis did not reveal major changes in G1, S, or G2/M distribution and neither in cell cycle progression (**Figures S2B-D**), but sh*Gen1* ESCs displayed a significant increase in cells with DNA content greater than 4C (**Figure 2F**). Metaphase spreads further showed a significant increase in chromosome fusions (**Figures 2G and 2H**), while chromosome breaks and micronuclei did not significantly change (**Figures 2G, 2I and S2F–H**).

Taken together, these data indicate that GEN1 depletion in ESCs is linked to genome destabilization characterized by accumulation of DNA content and structural chromosome fusions, in the absence of overt cell-cycle arrest. Within the limits of these assays, GEN1 thus emerges as an important contributor to chromosomal integrity in mouse ESCs.

### Catalytically inactive GEN1 rescues ESC self-renewal

To assess whether the requirement for GEN1 in ESC self-renewal depends on its nuclease activity, we generated *Gen1*^3xFL^ KI ESC lines stably overexpressing human GEN1 wild-type (hGEN1 WT), a catalytically inactive D157A mutant (hGEN1 MUT) (Ip *et al*., 2008), or an empty vector (EV) control (**Figures 3A, S3A and S3B**). We then performed CRISPR/Cas9-mediated disruption of endogenous 3×FLAG-tagged mouse *Gen1* in each background, followed by colony formation assays (**Figure 3B**). In EV-expressing ESCs, sgRNAs targeting *Gen1* markedly reduced AP^+^ colonies (**Figure 3C and 3D**), consistent with our previous loss-of-function results. Residual endogenous mouse GEN1^3xFL^ protein signal in a subset of surviving colonies suggested incomplete biallelic disruption (**Figure S3C**).

**Figure 3.**
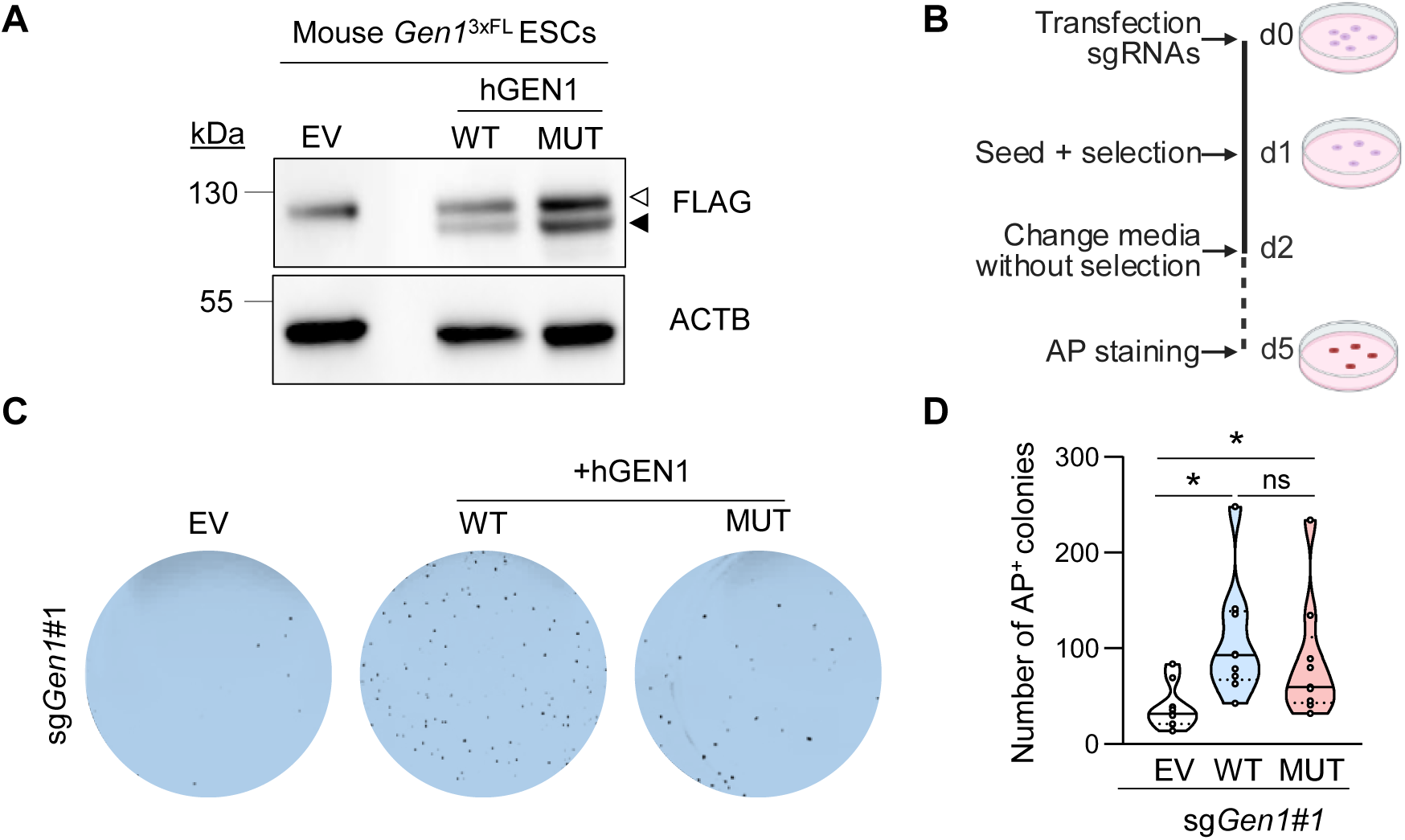
Catalytically inactive GEN1 is sufficient to rescue ESC colony formation. (A) Western blot showing expression of endogenous mouse GEN1 (upper band, white arrowhead) and exogenously expressed wild-type (WT) or catalytic mutant (MUT), human GEN1 (lower band, black arrowhead), detected with an anti-FLAG antibody. ACTB was used as loading control. (B) Schematic representation of the experimental procedure for CRISPR/Cas9-mediated KO of *Gen1* in ESCs and subsequent CFA. (C) Representative images of CFA wells from ESCs transfected with sg*Gen*1#1, with or without overexpression of WT or MUT hGEN1. (D) Quantification of AP+ colonies in *Gen1* KO ESCs with or without overexpression of hGEN1 WT or MUT. Data represent mean ± SEM (n = 8 biological replicates). Statistical significance was assessed by one-way ANOVA with Dunnett’s comparison test to EV conditions. \**p* < 0.05; ns, non-significant.

By contrast, both hGEN1 WT- and hGEN1 MUT-expressing ESCs maintained AP^+^ colony numbers upon endogenous *Gen1* disruption, whereas EV ESCs showed reduced AP^+^ colonies under the same conditions (**Figure 3C and 3D**). In the rescue backgrounds, endogenous mouse GEN1 was undetectable in surviving clones by immunoblotting (**Figure S3D and S3E**), indicating that expression of either WT or catalytically inactive human GEN1 is sufficient to sustain ESC colony formation in the absence of detectable endogenous GEN1 protein. These findings support that GEN1 nuclease activity is dispensable for ESC colony-forming capacity and suggest that non-catalytic functions of GEN1 make an important contribution to ESC maintenance.

### GEN1 influences ESC responses to topoisomerase I-mediated replication stress

Given the requirement for GEN1 in ESC maintenance, we next examined whether *Gen1* silencing alters ESC sensitivity to DNA damaging agents that generate distinct lesions (**Figure 4A**). MTT viability assays revealed no major differences between sh*Gen1* and sh*Luci* ESCs in response to cisplatin, bleomycin, olaparib, hydroxyurea (HU), or 4-nitroquinoline 1-oxide (4-NQO) (**Figures S4A-E**). In contrast, sh*Gen1* ESCs exhibited enhanced survival under camptothecin (CPT), a topoisomerase I (TOP1) poison (**Figure 4B**). To assess whether this effect reflects a general change in response to topoisomerase-mediated damage, we compared CPT with etoposide (ETOP), a topoisomerase II (TOP2) poison. Etoposide exposure did not produce a differential response between sh*Gen1* and sh*Luci* ESCs (**Figure 4C**), suggesting that GEN1 loss preferentially affects responses to TOP1-mediated replication stress.

**Figure 4.**
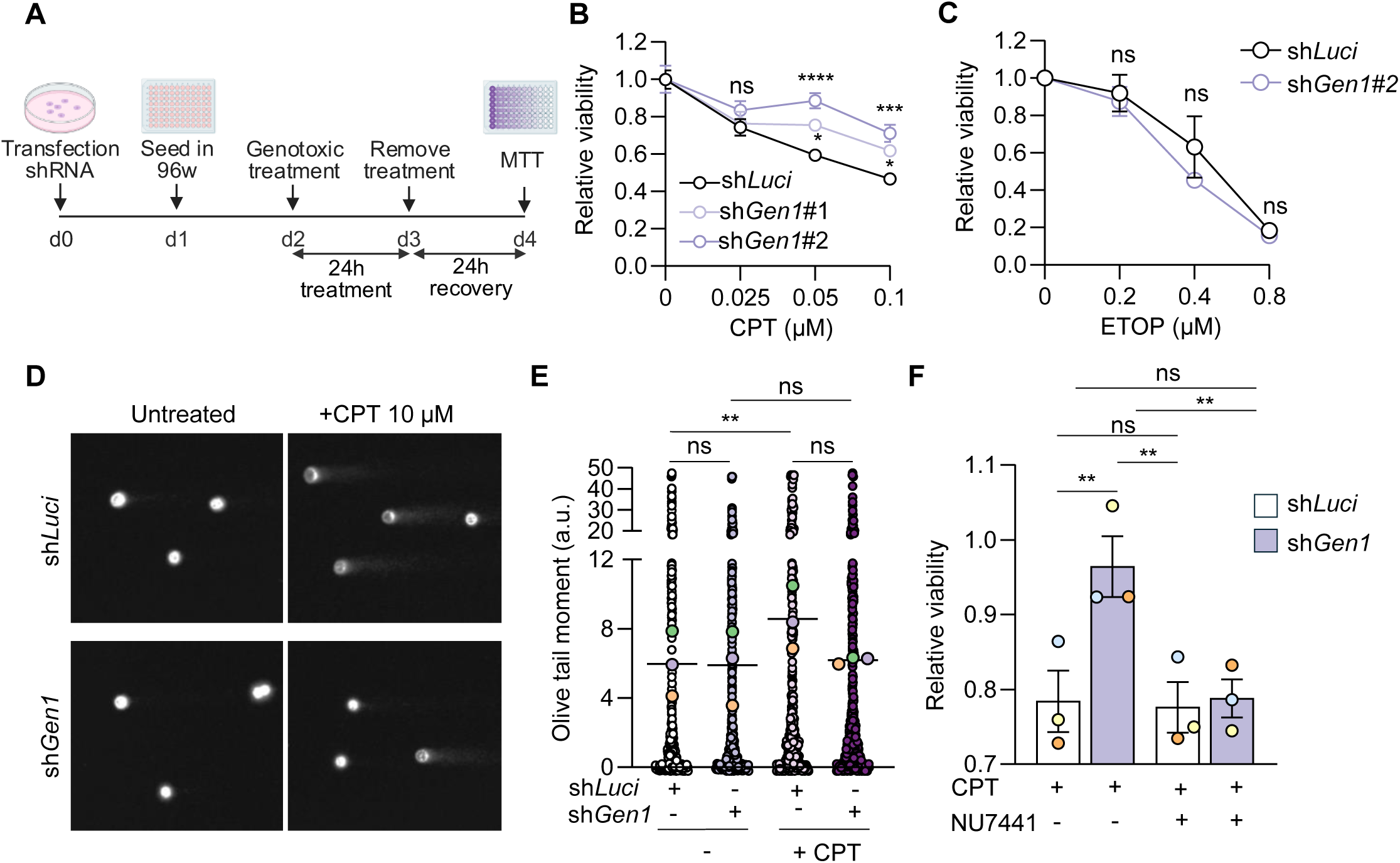
*Gen1* silencing affects the response of ESCs to camptothecin. (A) Schematic of the experimental approach for the MTT cell viability assay using camptothecin (CPT) and etoposide (ETOP). Survival curves of sh*Gen1* and sh*Luci* ESCs treated with increasing concentrations of CPT (B) or Etoposide (ETOP) (C) relative to untreated samples at 0 µM. Data represent mean ± SEM. (D) Representative comet assay images for each experimental condition. (E) Quantification of Olive tail moment (arbitrary units, a.u.) with or without 10 μM CPT treatment in *Gen1*- or *Luci*-silenced ESCs. (F) Relative viability to CPT-untreated samples of sh*Gen1* or sh*Luci* ESCs treated with 0.1 µM CPT in the presence or absence of the DNA-PKcs inhibitor NU7441 (10 µM). Statistical significance was assessed by two-way ANOVA with Dunnett’s comparisons test (B) or Sidak comparison test (C), one-way ANOVA with Tukey’s multiple comparisons test (E) or two-way ANOVA with Tukey’s multiple comparisons test (F). \**p* < 0.05; \*\**p* < 0.01; *** *p* < 0.001; **** *p* <0.0001; ns, non-significant. Data represent mean ± SEM.

Consistent with this finding, alkaline comet assays showed reduced Olive tail moments in sh*Gen1* ESCs compared with controls after CPT exposure, indicating decreased DNA breakage under our assay conditions (**Figures 4D and 4E**). These results are consistent with altered processing of CPT-stabilized TOP1-DNA cleavage complexes when GEN1 is depleted, rather than simply an overall increase in resistance to DNA damage. Taken together, these data indicate that GEN1 contributes to the ESC response to topoisomerase I poisoning. The unexpected increase in CPT tolerance upon *Gen1* silencing led us to hypothesize that ESCs might adjust the repair pathway usage to maintain short-term viability. To explore this possibility, we performed viability assays in the presence or absence of the DNA-PK inhibitor NU7441, which impairs classical non-homologous end joining (NHEJ) (Leahy *et al*., 2004). In sh*Gen1* ESCs, NU7441 treatment abolished the CPT-associated survival advantage, restoring viability to levels comparable to sh*Luci* controls, whereas NU7441 had minimal impact on CPT responses in sh*Luci* ESCs (**Figure 4F**). Within the limits of these assays, this pattern is compatible with increased reliance on DNA-PK-dependent NHEJ in GEN1-deficient ESCs under TOP1-induced replication stress, although direct measurements of HR and NHEJ activity will be required to determine how repair pathway choice is altered at the mechanistic level.

## Discussion

GEN1 emerges as indispensable for ESC self-renewal in a manner that distinguish its pluripotent function from roles in somatic cells. Unlike its reported backup role behind MUS81 and BLM dissolution in somatic contexts (Wang *et al*., 2016; Garner *et al*., 2013), GEN1 proves non-redundant in ESCs, where its depletion abolishes colony formation, a phenotype specifically suppressed by expression of the human GEN1 ortholog. This ESC-specific requirement may arise from the distinctive cell cycle features of ESC, including a shortened G1 phase and elevated replication stress associated with their highly proliferative profile (White *et al*., 2005; Ahuja *et al*., 2016).

Consistent with an essential role in safeguarding genome integrity, GEN1 depletion compromises ESC self-renewal while leaving core pluripotency markers largely unchanged. Rather than triggering a classical cell-cycle arrest, *Gen1* knockdown leads to an accumulation of cells with more than 4C DNA content and a pronounced increase in structural chromosome aberrations, most notably chromosome fusions, without a rise in chromosome breaks or micronuclei. These phenotypes are compatible with unresolved replication or recombination intermediates persisting into mitosis and being converted into structural rearrangements, in line with previous work linking unresolved joint molecules, arising from loss of *Gen1*, to chromosomal aberrations (Chan, Fugger and West, 2018). Together, these observations indicate that GEN1 loss drives genome instability in ESCs primarily through accumulation of DNA and fusion events rather than overt checkpoint activation or loss of pluripotency.

Unexpectedly, the essential role of GEN1 in ESC maintenance *in vitro* is independent of its catalytic activity. A catalytically inactive hGEN1 D157A mutant (Rass *et al*., 2010; Chan and West, 2015) rescues colony formation in *Gen1*-null ESCs, uncoupling viability from nuclease function. This result indicates that nuclease activity is not required for all GEN1-dependent functions in ESCs and suggests that catalytic-independent activities may contribute substantially to the maintenance of pluripotent cells. Recent work in several model organisms reporting DNA repair-independent functions of GEN1 further supports the notion that this protein contributes to genome stability through both catalytic and non-catalytic mechanisms (Bailly *et al*., 2010; Chen and Aström, 2012; Budzyk *et al*., 2025). At present, however, the underlying mechanism remains unresolved. GEN1 may help stabilize repair or replication-associated assemblies, influence the handling of DNA intermediates independently of cleavage, or participate indirectly in pathways that preserve proliferative fitness.

Moreover, non-canonical roles of GEN1 might extend to replication stress coping mechanisms. The response to camptothecin (CPT) adds another layer to this interpretation. GEN1-depleted ESCs showed increased short-term tolerance to TOP1 poisoning, reduced comet tail moments under CPT treatment, and selective sensitivity to DNA-PK inhibition in this setting. Taken together, these observations suggest that loss of GEN1 changes how ESCs process TOP1-associated replication lesions and may favour repair routes that depend more strongly on DNA-PK activity. Nevertheless, the present experiments do not directly measure HR or NHEJ flux, and therefore the proposal that GEN1 influences double-strand break repair pathway usage should be viewed as a working model supported indirectly by pharmacological and survival data.

From a broader pluripotent stem cell perspective, the genomic abnormalities arising upon GEN1 loss, including structural chromosome fusions and altered responses to replication stress, resemble classes of defects that are of concern during prolonged pluripotent stem cell culture (Mayshar *et al*., 2010; Spits *et al*., 2008; Lefort *et al*., 2008). In this context, GEN1 emerges as a relevant component of the machinery that helps maintain genomic stability in ESCs. Extending these observations to human PSCs and testing directly how GEN1 influences repair pathway usage will be important next steps for evaluating its significance in stem cell quality control and long-term culture fitness.

## Resources availability

### Lead contact

Further information and requests for resources and reagents should be directed to and will be fulfilled by the lead contact, Diana Guallar.

### Materials availability

The knock-in *Gen1^3xFl^*ESC line generated in this study is available upon request and subject to an MTA.

### Data and code availability

Any additional information required to reanalyze the data reported in this paper is available from the lead contact upon request.

## Acknowledgements

This work was supported by grants from MICIU/AEI/10.13039/501100011033/FEDER, UE (D.G.: PID2022-136608OB-I00, M.G.B: PID2023-147024NB-I00, M.F.: PID2022-143105NB-I00), Xunta de Galicia (D:G: ED431C2023/28, ED431C 2023/28 and ED481A-2023-138; M.G.B.: ED431C 2023/10) and FEDER ‘Una manera de hacer Europa’ (D.G.: ED431C-2023/28). D.G. was a recipient of Ramón y Cajal (RYC2019-027305-I) award from the Ministerio de Ciencia e Innovación. L.R-L. (Xunta de Galicia, ED481A-2023-138), D.M.-D. (Xunta de Galicia, ED481A-2023-026), T.M. (Fundação para a Ciência e Tecnologia of Portugal, 2021.07526.BD), V.G.-O.(Ministerio de Ciencia, Innovación y Universidades, FPU17/01131) and A.C. (Xunta de Galicia, ED481A 2021/053) were beneficiaries of fellowships. L.R.-L. was a beneficiary of an EMBO Scientific Exchange Grant (11185).

## Author contributions

Conceptualization, M.G.B. and D.G.; methodology, L.R.-L., C.A., D.M.-D., H.C.-M., T.M., R.C., D.R.-C., A.C., A.F.-I., D.G. and M.G.B.; formal analysis, L.R.-L. and D.G.; investigation, L.R.-L., C.A., H.C.-M., T.M., R.C., D.R.-C., E.S. and M.F.; writing - original draft preparation, L.R.-L., M.F., M.G.-B. and D.G.; writing - review and editing, L.R.-L., C.A., D.M.-D., T.M., R.C., D.R.-C., A.F.-I., E.S., M.F., M.G.B. and D.G.; funding acquisition, E.S., M.F., M.G.B. and D.G.; supervision, D.G.

## Declaration of generative ai and ai-assisted technologies in the writing process

During the preparation of this work, the authors used Claude AI to enhance the clarity of the text. The authors reviewed and edited the content and take full responsibility.

## STAR Methods

### Key Resources Table

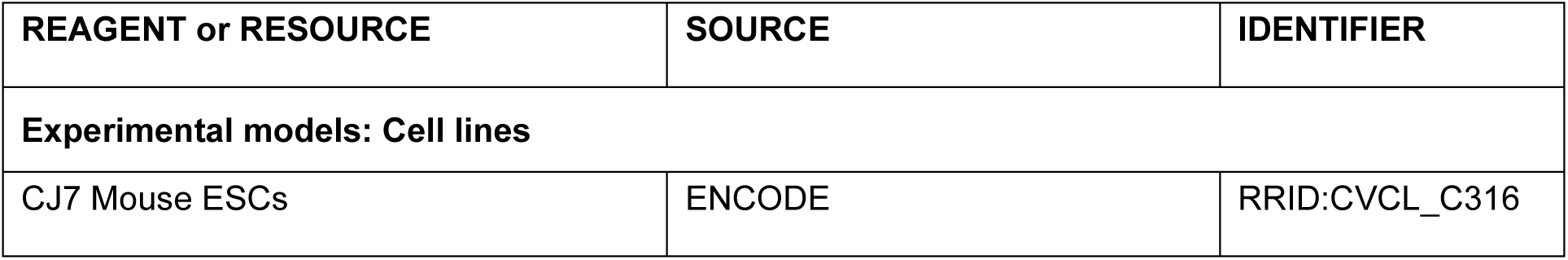
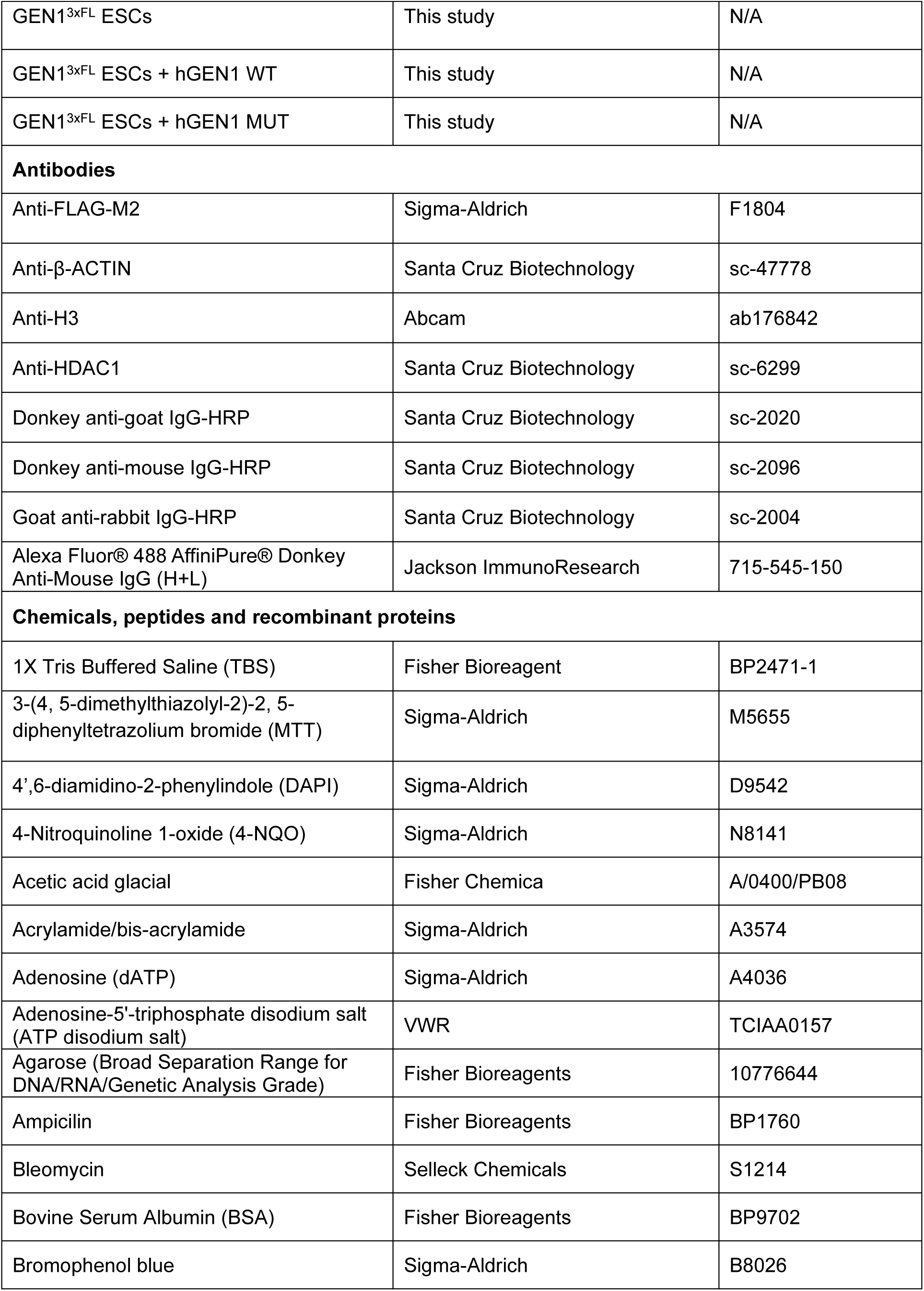
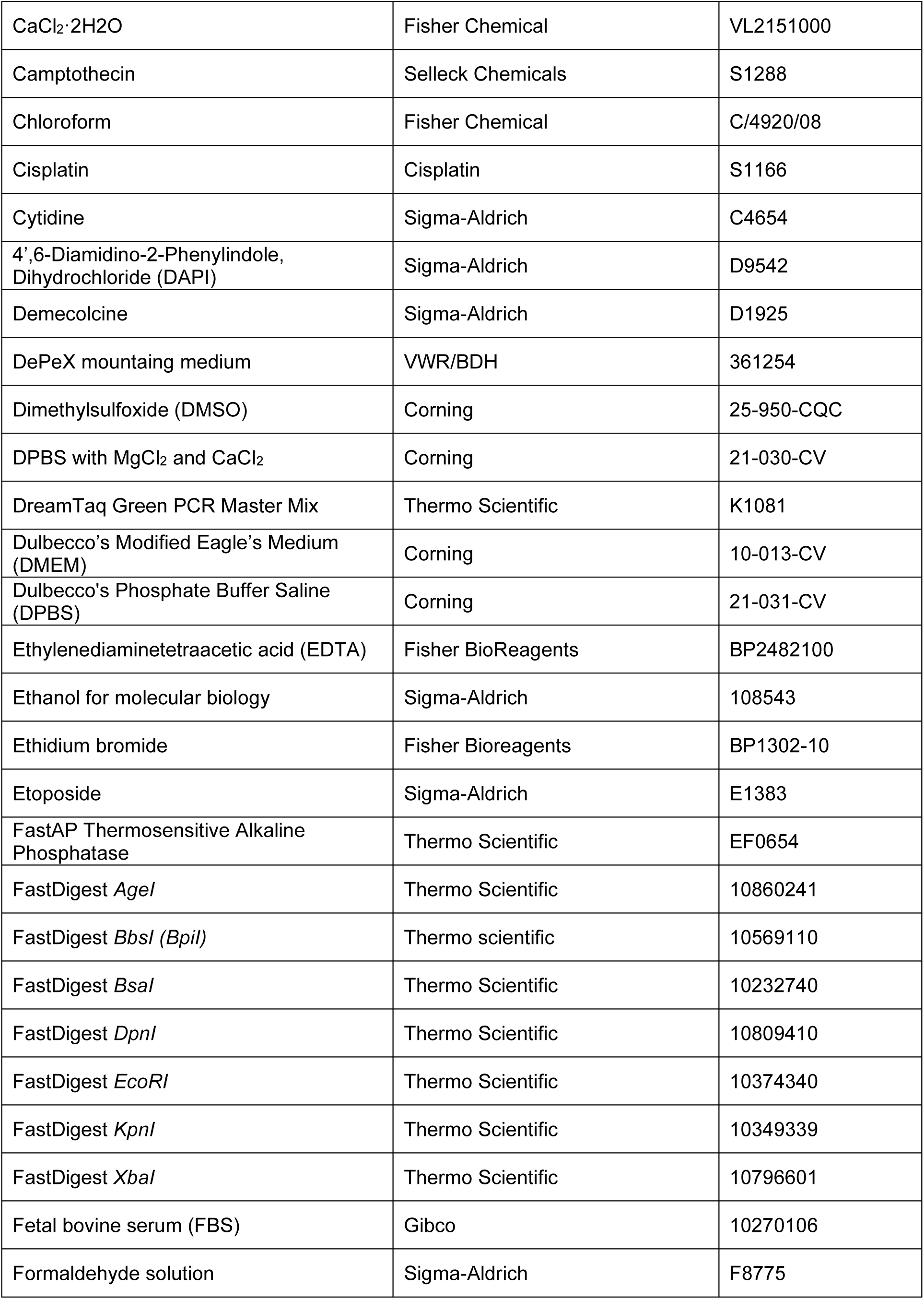
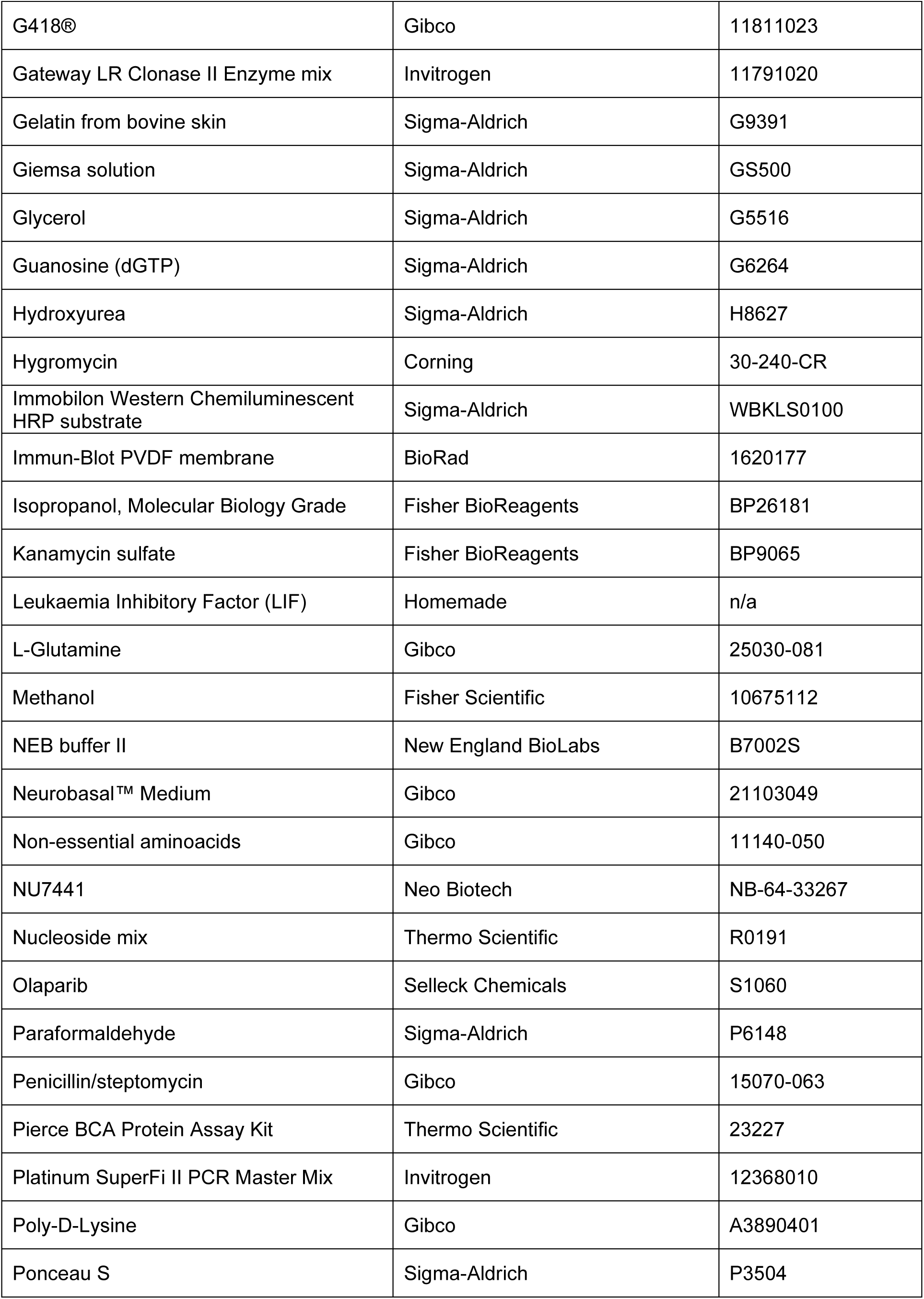
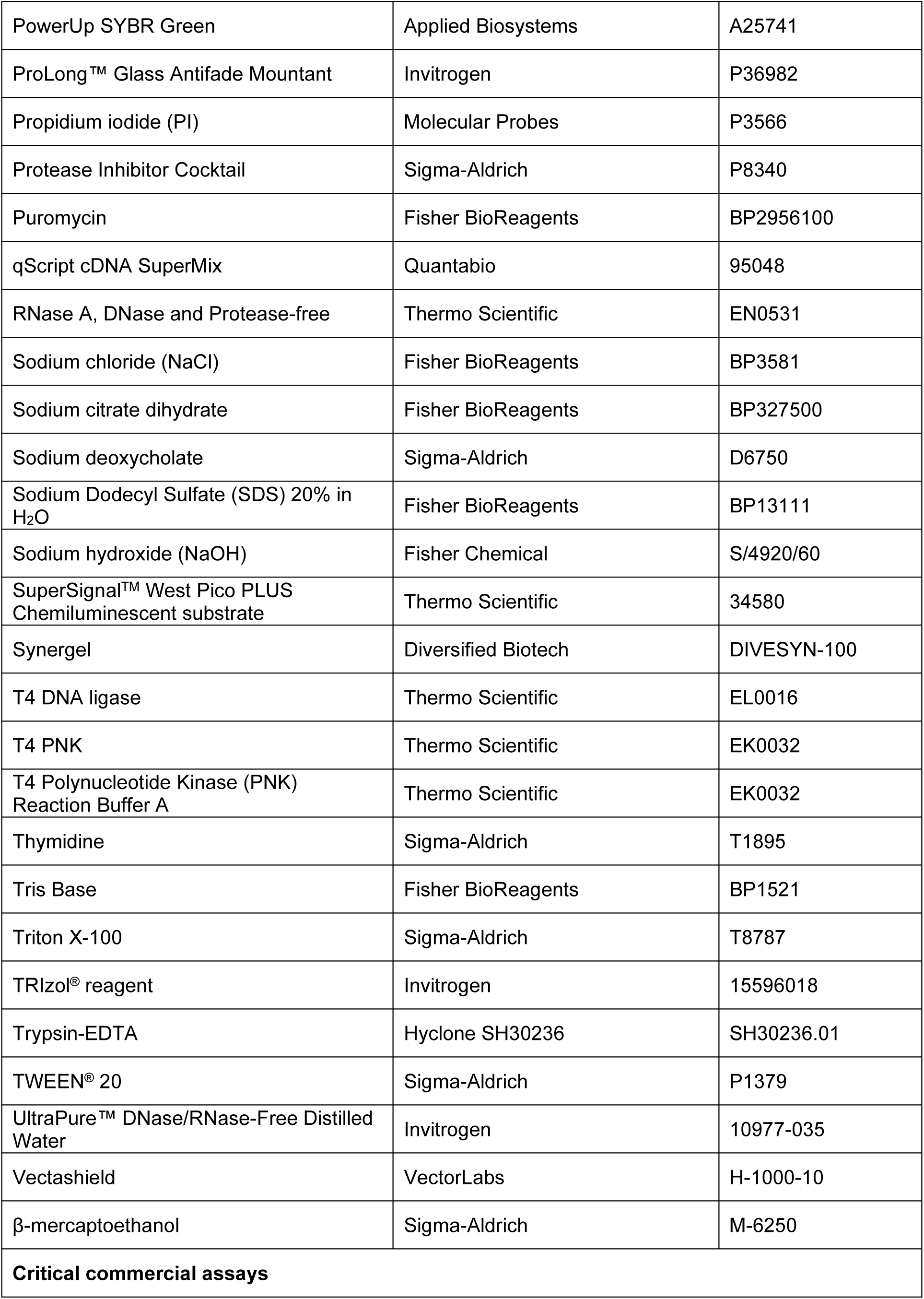
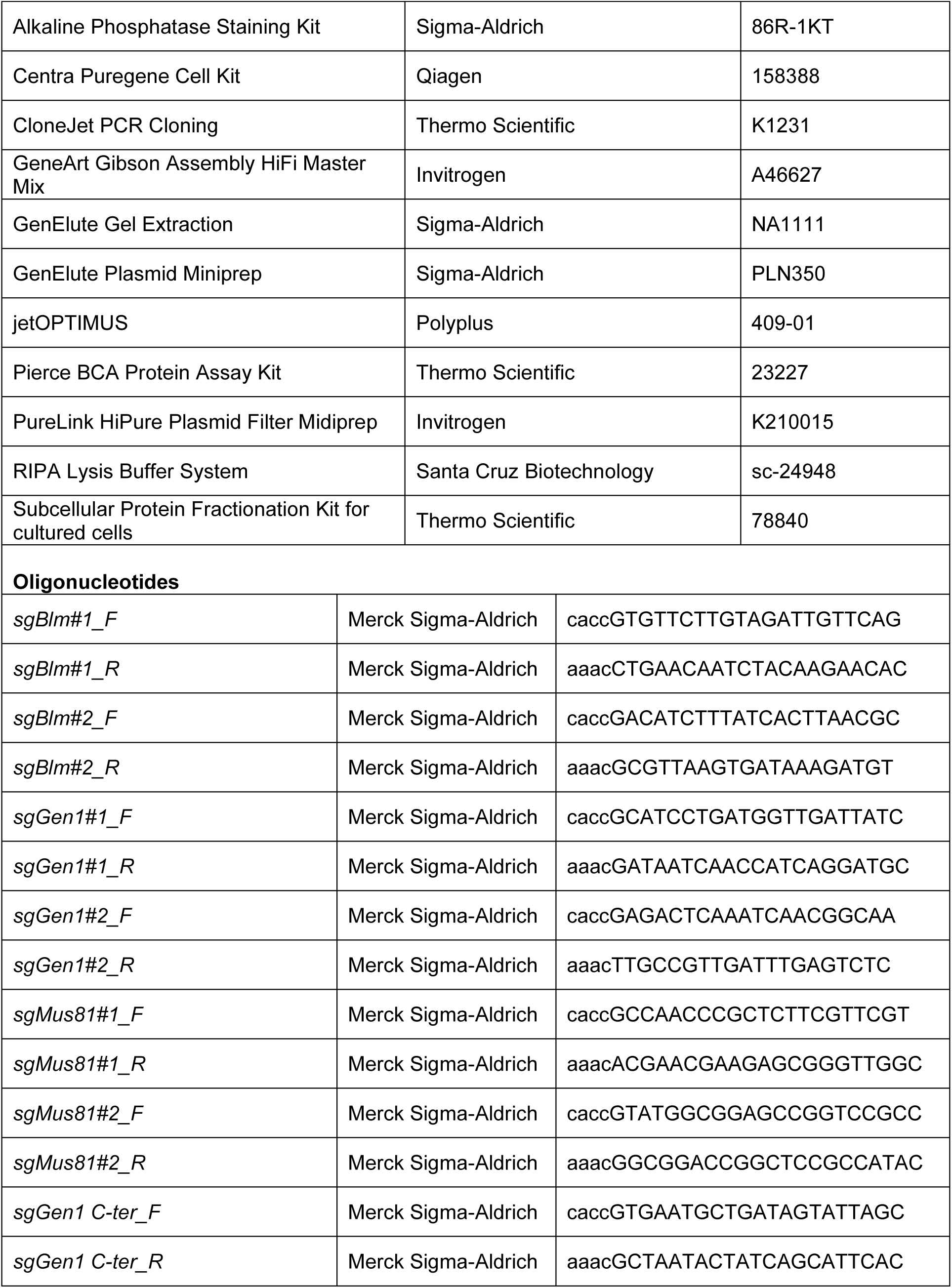
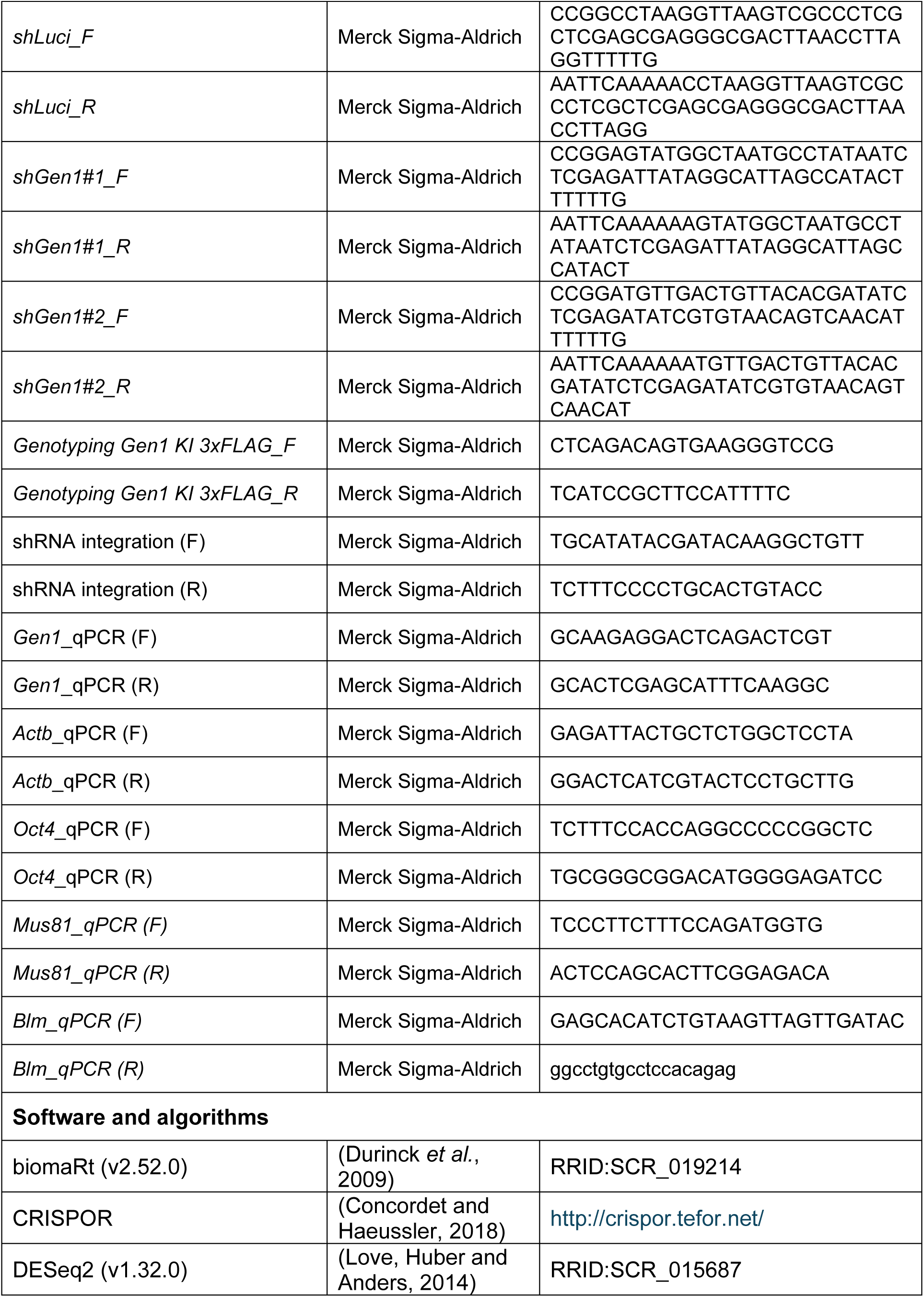
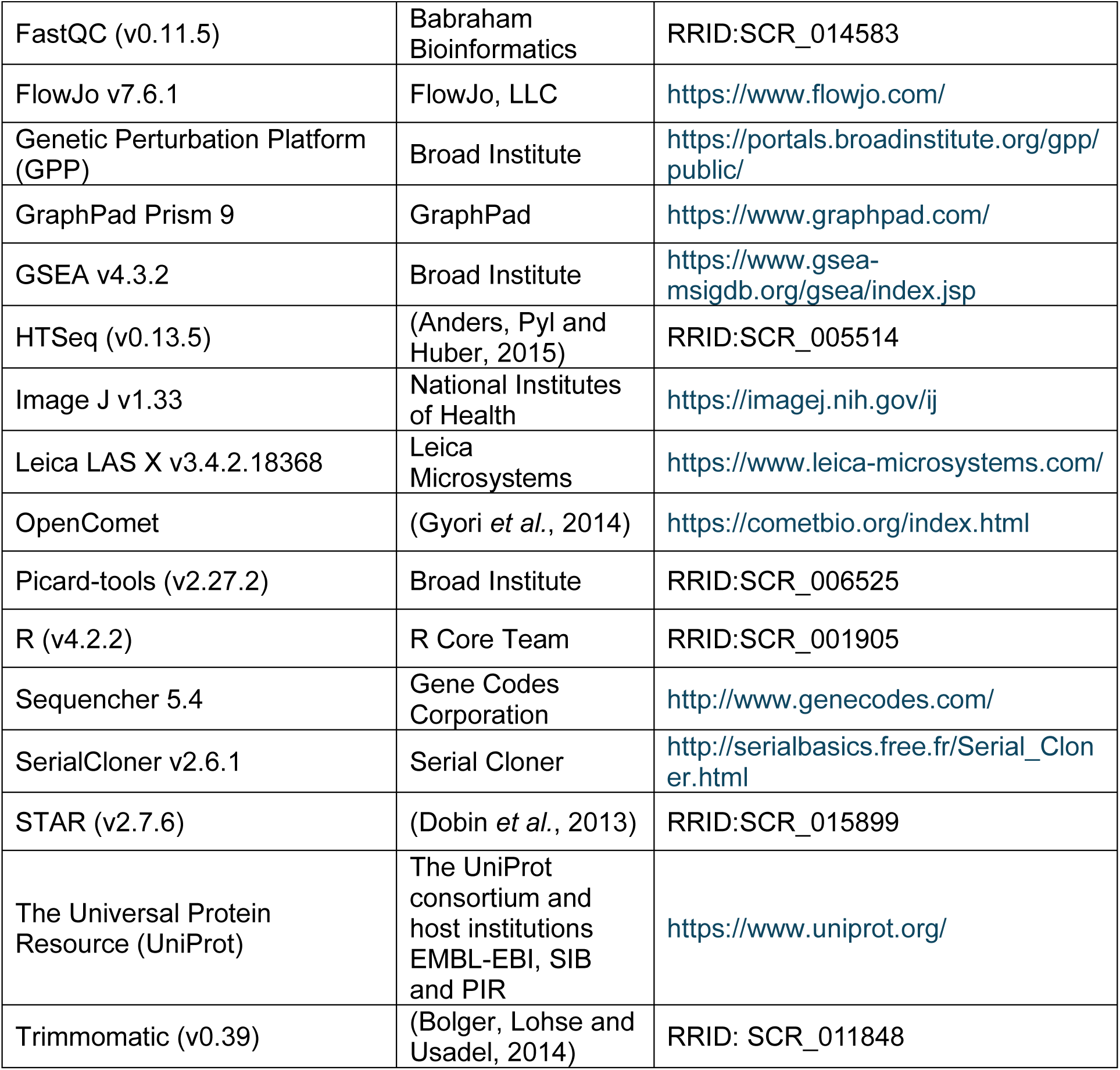

## Contact for Reagent and Resource Sharing

Further information and requests for resources should be directed to and will be fulfilled by the lead contact, Diana Guallar.

## Experimental Model and Study Participant Details

This study used established mouse ESC lines and did not involve living animals or human participants. The wild-type CJ7 mouse ESC line served as the parental line for all genetic modifications. We generated and analysed CRISPR/Cas9-edited GEN1^3xFL^ homozygous knock-in ESCs, as well as stable lines overexpressing human GEN1 WT or catalytically inactive D157A mutant alongside empty vector controls. All ESC lines were derived from the male CJ7 background and authenticated by genotyping PCR and anti-FLAG immunoblotting. The experimental design did not test for sex-specific effects, as all lines share the same genetic background; therefore, findings represent outcomes specific to these male-derived ESC lines. All procedures involving mouse ESCs were conducted in accordance with institutional biosafety guidelines.

## Method Details

### Mouse ESC culture

Mouse embryonic stem cells (ESCs) were cultured on gelatin-coated plates prepared by incubating plates with 0.1% gelatin (Sigma-Aldrich) in Milli-Q water for at least 30 min at 37°C. ESCs were maintained in ESM^LIF^ medium consisting of high-glucose DMEM (Corning), 15% heat-inactivated fetal bovine serum (FBS) (Corning), 100 U/mL penicillin/streptomycin (Gibco, 15140-122), 1% nucleoside mix (adenosine [Sigma-Aldrich, 10 μg/mL], cytidine [Sigma-Aldrich; 10 μg/mL], guanosine [Sigma-Aldrich; 10 μg/mL], thymidine [Sigma-Aldrich; 10 μg/mL], uridine [Sigma-Aldrich; 10 μg/mL]), 2 mM L-glutamine (Gibco), 1× non-essential amino acids (Gibco), 0.1 mM β-mercaptoethanol (Sigma-Aldrich), and 1,000 U/mL LIF. Cells were passaged every other day at a 1:8 ratio using 0.05% trypsin-EDTA (Hyclone) for 7 min at 37°C and maintained at 37°C in 5% CO2.

### Transient transfection

ESCs were seeded at 15,000 cells/cm^2^ the day before transfection to reach 60%–80% confluence. Transfections were performed using jetOPTIMUS (Polyplus) according to the manufacturer’s instructions. Briefly, for a well of 6-well plate, 2 μg of DNA were diluted in 200 μL jetOPTIMUS buffer, mixed with 2 μL reagent, incubated for 10 min at room temperature, and added dropwise to the cells. The following day, cells were trypsinised and transferred to antibiotic selection medium, which was replaced daily until untransfected control cells were dead.

### Colony formation assay

For colony formation assays, 800 cells per well were seeded in 6-well gelatin-coated plates or 400 cells per well in 12-well plates. Medium was changed daily for 4–5 days. Colonies were fixed and stained using the alkaline phosphatase (AP) staining kit (Sigma-Aldrich). For fixation, a solution containing 0.1 M sodium citrate, acetone, and 37% formaldehyde was applied for 30 s at room temperature, followed by rinsing with deionized water. Staining was performed using sodium nitrite, FRV-alkaline, and Naphtol AS-BI alkaline according to the manufacturer’s instructions. AP-positive (AP+) colonies were scored by phase-contrast microscopy.

### CRISPR/Cas9-mediated knock-in and genotyping

WT ESCs were transfected with an sgRNA targeting the 3’ region of *Gen1* coding sequence together with a donor plasmid encoding a 3xFLAG-P2A-NeoR cassette flanked by *Gen1* 3′ homology arms. Cells were selected with G418 sulfate (Gibco) until untransfected controls were dead. Individual colonies were isolated and analysed by DNA amplification and protein expression. For knock-in genotyping, PCR primers flanking the insertion site were designed to amplify amplicons of 1,619 bp for the WT allele and 2,598 bp for the KI allele.

### shRNA cloning and knockdown

shRNAs were designed using the Broad GPP portal and selected for UTR targeting and high efficiency scores. Oligonucleotides were annealed by mixing 2.5 μM of each oligo with 1× NEB buffer 2 (New England Biolabs), heating to 95°C for 10 min, 70°C for 10 min, and cooling at a rate of −1°C per 2.5 min to 25°C before holding at 4°C. Annealed oligos were cloned into pHSVg vectors digested with *Age*I and *Eco*RI (Thermo Scientific), gel purified, ligated with T4 DNA ligase (Thermo Scientific), and transformed into Stbl3 cells. Constructs were confirmed by colony PCR and Sanger sequencing.

### RNA Extraction and Quantitative PCR (RT-qPCR)

RNA was extracted using either the E.Z.N.A. Total RNA Kit I (Omega Bio-Tek) or TRIzol reagent (Invitrogen) following the respective manufacturer’s instructions. For TRIzol-based extraction, cells were lysed in 600 μL of TRIzol per well of a 6-well plate, scraped, and transferred to Phasemaker tubes (Invitrogen) following chloroform phase separation (Fisher Scientific). RNA was precipitated with isopropanol (Fisher Scientific), washed with ice-cold ethanol (Sigma-Aldrich), air-dried, and dissolved in 20–50 μL of UltraPure DNase/RNase-free distilled water (Invitrogen). RNA concentration and purity were assessed using a NanoDrop 2000 spectrophotometer (Thermo Scientific). Total RNA (1 μg) was converted into cDNA using the qScript cDNA SuperMix (Quantabio) following the manufacturer’s instructions. Quantitative PCR was performed using the PowerUp SYBR Green Master Mix (Applied Biosystems) with gene-specific primers on a QuantStudio 5 real-time PCR system (Applied Biosystems). Data were analysed using the delta-delta Ct (ΔΔCt) method (Livak and Schmittgen, 2001), normalised to the housekeeping gene *Actb*, and expressed as relative expression to the indicated control condition.

### Transcriptomic analysis by RNA-sequencing

For total RNA sequencing, ESCs transfected with sh*Gen1* or sh*Luci* as control were lysed with TRIzol and total RNA was obtained following manufactureŕs instructions. RNA quality was verified using an Agilent Bioanalyzer 2100 by NOVOGENE, followed by library construction with Ribo-Zero® kit (Illumina, 20040525). RNA-seq libraries were sequenced on a HiSeq 4000 platform using a 150 bp paired-end, unstranded protocol.

For differential gene expression analysis, sequencing adaptors were removed using the software Trimmomatic (v0.39) and the quality of the reads was assessed using FastQC (v0.11.5) software. For read alignment to the mouse genome (Ensembl GRCm39) and removal of sequencing duplicates, the software packages STAR (v2.7.6) and Picard-tools (v2.27.2) were used. After the filtering, reads were converted into gene counts using HTSeq (v0.13.5). Gene counts analysis was performed using R (v4.2.2) with the DESeq2 (v1.32.0) package. The resulting data were annotated with gene ID, chromosome, start and end position, strand and transcript type with the BiomaRT (v2.52.0) package.

### Metaphase spreads

Cells were treated with 0.2 μg/mL demecolcine (Sigma-Aldrich) for 2 h at 37°C. Cells and media were collected, pelleted, and resuspended in PBS. Hypotonic treatment was performed with 0.03 M trisodium citrate solution prewarmed to 37°C for 30 min. Cells were then fixed three times in methanol:acetic acid (3:1), dropped onto acetic acid-pretreated slides, and air-dried. Slides were stained with Giemsa solution (Sigma-Aldrich, 1:20 in water) for 20 min, rinsed, and dried overnight. Mounted slides were imaged using a Leica DM4B or Zeiss AxioObserver Z1 microscope, and 50 metaphases per condition were scored.

### Alkaline comet assay

For alkaline comet assays, 20,000 cells per condition were pelleted, resuspended in 0.5% low-melt agarose in PBS, and applied to 1.5% agarose-coated slides. After solidification, slides were lysed for 1 h at 4°C in alkaline lysis buffer. Electrophoresis was performed after a 30 min equilibration in 300 mM NaOH/1 mM EDTA at 25 V for 25 min. Slides were neutralized in 0.4 M Tris-HCl (pH 7.5), dehydrated in 100% ethanol, air-dried, and stained with 1 μM DAPI. At least 100 comets per condition were scored using OpenComet in ImageJ, and Olive tail moment pluggin was used for quantification. As a positive control for DNA breakage, cells were treated with 10µM CPT for 4 hours.

### MTT viability assay

ESCs were seeded at 5,000 cells per well in 96-well plates. The following day, cells were exposed to genotoxic agents for 24 h and then returned to drug-free ESM^LIF^ for an additional 24 h. MTT (Sigma-Aldrich, 0.5 mg/mL) was added for 4 h at 37°C in the dark. Formazan crystals were solubilized in isopropanol:DMSO (1:1) with shaking for 15 min at room temperature, and absorbance was measured at 570 nm with background correction at 690 nm. Viability was normalized to vehicle-treated controls.

### Immunofluorescence

12,000 cells were seeded in wells of 48-well plates coated with 0.1% gelatin and cultured in ESCs standard medium. After 24 h, cells were fixed with 4% paraformaldehyde for 15 min in darkness at RT and washed two times with PBS (Dulbecco’s Phosphate Buffered Saline, Sigma-Aldrich). Then, cells were permeabilizated with 0.25% Triton X-100 (Sigma-Aldrich) diluted in PBS for 5 min at RT followed by two times of PBS washes and blocking with 10% BSA (Fisher Bioreagents) diluted in PBS for 30 min at 37°C. Cells were incubated with the primary antibody of choice diluted in 1% BSA in PBS for 1 h at RT in a humidity chamber. Cells were washed three times with 0.1% Tween in PBS for 5 min each time. Protected from light, cells were incubated with the appropriate secondary antibody in 1% BSA in PBS for 1 h at room temperature in a humidity chamber. Next, cells were washed three times with 0.1% Tween in PBS for 5 min each time. Then, they were washed once with PBS for 5 min followed by another wash with milli-Q water for another 5 min. The coverslips were air-dried completely at RT. Coverslips were mounted on the slides using ProLong™ Glass Antifade Mountant (Invitrogen). Coverslips were sealed with nail varnish and air-dried. Images were acquired with ZEISS Celldiscoverer 7 microscope using a 40x objective. Nuclei were stained with DAPI (4′,6-diamidino-2-phenilindole, Sigma-Aldrich).

### Micronuclei detection

Cells were seeded on coverslips previously coated with 50 μg/mL poly-D-Lysine (Gibco). The plate containing the coverslips was placed on ice and washed twice with PBS. Cells were fixed with 4% PFA for 10 min at RT. Cells were washed twice with PBS after fixation at RT. Cells were permeabilised with 0.2% Triton-X in PBS for 5 min and then washed twice with PBS. Cells were stained with 1 μg/mL DAPI (Sigma-Aldrich). Then, they were washed once with PBS for 5 min followed by another wash with milli-Q water for another 5 min. The coverslips were air-dried completely at RT. Coverslips were mounted on slides using Vectashield® (Vector Labs) mounting medium. They were sealed with nail varnish and air-dried. Images were acquired using a ZEISS LSM 880 confocal microscope. The number of micronuclei was scored after checking at least 500 cells per condition.

### Cell cycle analysis by flow cytometry

Cells were trypsinized, washed with PBS, and fixed by slow addition of ice-cold 70% ethanol with shaking for 30 min at 4°C. Fixed cells were washed twice with PBS and incubated in staining solution containing propidium iodide and RNase A for 30 min at 37°C in the dark. Samples were analyzed on a FACSCalibur cytometer, and data were processed using FlowJo with the Watson pragmatic model.

### G1/S synchronization

ESCs were seeded at 30,000 cells/cm^2^ and subjected to a double thymidine block. Cells were first treated with 2 mM thymidine (Sigma-Aldrich) for 16 h, released into fresh ESM^LIF^ for 8 h, and then blocked again with 2 mM thymidine for 16 h. Cells were released into fresh medium, and samples were collected at the indicated time points.

### Subcellular fractionation

Subcellular fractions were prepared using the Subcellular Protein Fractionation Kit (Thermo Scientific) according to the manufacturer’s protocol. Between 10^6^ and 10^7^ cells per condition were harvested and fractionated into cytoplasmic, membrane, nuclear soluble, chromatin-bound, and cytoskeletal fractions. Protein concentrations were determined by Pierce BCA protein assay (Thermo Scientific) using fraction-specific standard curves.

### Protein extraction

Whole-cell lysates were prepared in RIPA Lysis buffer system (Santa Cruz Biotechnology) which includes Protease Inhibitor Cocktail, PMSF and sodium orthovanadate. Cells were lysed on ice for 30 min with intermittent vortexing, cleared by centrifugation for 10 minutes at 10,000 x *g* at 4°C, and protein concentrations were determined by Pierce BCA protein assay (Thermo Scientific).

### Western blotting

Lysates were mixed with Laemmli buffer, denatured at 95°C for 5 min, and separated by SDS-PAGE on 8%-12% homemade gels or 4%-20% Tris-Glycine gels. Proteins were transferred to PVDF membranes using semi-dry or wet transfer conditions. Membranes were stained with Ponceau S, blocked in 5% skim milk in TBS-T, and incubated with primary antibodies in blocking buffer. After washing, membranes were incubated with HRP-conjugated secondary antibodies and developed using chemiluminescent substrate. Images were acquired on a ChemiDoc system.

### Statistical analysis

Statistical analyses were performed using GraphPad Prism or R. Details of the statistical tests used for each experiment are provided in the corresponding figure legends. Data are presented as mean ± SEM from at least three biological replicates. Independent biological replicates are colour coded. Differences were considered significant at *p-value* < 0.05.

## References

Ahuja, A. K., Jodkowska, K., Teloni, F., Bizard, A. H., Zellweger, R., Herrador, R., Ortega, S., Hickson, I. D., Altmeyer, M., Mendez, J. and Lopes, M. (2016) ‘A short G1 phase imposes constitutive replication stress and fork remodelling in mouse embryonic stem cells’, Nat Commun, 7, pp. 10660.

Anders, S., Pyl, P. T. and Huber, W. (2015) ‘HTSeq--a Python framework to work with high-throughput sequencing data’, Bioinformatics, 31(2), pp. 166–9.

Bailly, A. P., Freeman, A., Hall, J., Déclais, A. C., Alpi, A., Lilley, D. M., Ahmed, S. and Gartner, A. (2010) ‘The Caenorhabditis elegans homolog of Gen1/Yen1 resolvases links DNA damage signaling to DNA double-strand break repair’, PLoS Genet, 6(7), pp. e1001025.

Banáth, J. P., Bañuelos, C. A., Klokov, D., MacPhail, S. M., Lansdorp, P. M. and Olive, P. L. (2009) ‘Explanation for excessive DNA single-strand breaks and endogenous repair foci in pluripotent mouse embryonic stem cells’, Exp Cell Res, 315(8), pp. 1505–20.

Bolger, A. M., Lohse, M. and Usadel, B. (2014) ‘Trimmomatic: a flexible trimmer for Illumina sequence data’, Bioinformatics, 30(15), pp. 2114–20.

Budzyk, M., Simon, A., Mace, A. S. and Basto, R. (2025) ‘A novel DNA repair-independent role for Gen nuclease in promoting unscheduled polyploidy cell proliferation’, PLoS Genet, 21(9), pp. e1011605.

Chan, Y. W., Fugger, K. and West, S. C. (2018) ‘Unresolved recombination intermediates lead to ultra-fine anaphase bridges, chromosome breaks and aberrations’, Nat Cell Biol, 20(1), pp. 92–103.

Chan, Y. W. and West, S. (2015) ‘GEN1 promotes Holliday junction resolution by a coordinated nick and counter-nick mechanism’, Nucleic Acids Res, 43(22), pp. 10882-92.

Chan, Y. W. and West, S. C. (2014) ‘Spatial control of the GEN1 Holliday junction resolvase ensures genome stability’, Nat Commun, 5, pp. 4844.

Chen, J. and Aström, S. U. (2012) ‘A catalytic and non-catalytic role for the Yen1 nuclease in maintaining genome integrity in Kluyveromyces lactis’, DNA Repair (Amst), 11(10), pp. 833–43.

Concordet, J. P. and Haeussler, M. (2018) ‘CRISPOR: intuitive guide selection for CRISPR/Cas9 genome editing experiments and screens’, Nucleic Acids Res, 46(W1), pp. W242–W245.

Dehé, P. M. and Gaillard, P. H. L. (2017) ‘Control of structure-specific endonucleases to maintain genome stability’, Nat Rev Mol Cell Biol, 18(5), pp. 315–330.

Dobin, A., Davis, C. A., Schlesinger, F., Drenkow, J., Zaleski, C., Jha, S., Batut, P., Chaisson, M. and Gingeras, T. R. (2013) ‘STAR: ultrafast universal RNA-seq aligner’, Bioinformatics, 29(1), pp. 15–21.

Durinck, S., Spellman, P. T., Birney, E. and Huber, W. (2009) ‘Mapping identifiers for the integration of genomic datasets with the R/Bioconductor package biomaRt’, Nat Protoc, 4(8), pp. 1184–91.

Fujii-Yamamoto, H., Kim, J. M., Arai, K. and Masai, H. (2005) ‘Cell cycle and developmental regulations of replication factors in mouse embryonic stem cells’, J Biol Chem, 280(13), pp. 12976–87.

Garner, E., Kim, Y., Lach, F. P., Kottemann, M. C. and Smogorzewska, A. (2013) ‘Human GEN1 and the SLX4-associated nucleases MUS81 and SLX1 are essential for the resolution of replication-induced Holliday junctions’, Cell Rep, 5(1), pp. 207–15.

Gowen, L. C., Johnson, B. L., Latour, A. M., Sulik, K. K. and Koller, B. H. (1996) ‘Brca1 deficiency results in early embryonic lethality characterized by neuroepithelial abnormalities’, Nat Genet, 12(2), pp. 191–4.

Gyori, B. M., Venkatachalam, G., Thiagarajan, P. S., Hsu, D. and Clement, M. V. (2014) ‘OpenComet: an automated tool for comet assay image analysis’, Redox Biol, 2, pp. 457–65.

Hakem, R., de la Pompa, J. L. and Mak, T. W. (1998) ‘Developmental studies of Brca1 and Brca2 knock-out mice’, J Mammary Gland Biol Neoplasia, 3(4), pp. 431–45.

Hakem, R., de la Pompa, J. L., Sirard, C., Mo, R., Woo, M., Hakem, A., Wakeham, A., Potter, J., Reitmair, A., Billia, F., Firpo, E., Hui, C. C., Roberts, J., Rossant, J. and Mak, T. W. (1996) ‘The tumor suppressor gene Brca1 is required for embryonic cellular proliferation in the mouse’, Cell, 85(7), pp. 1009–23.

Hanada, K., Budzowska, M., Modesti, M., Maas, A., Wyman, C., Essers, J. and Kanaar, R. (2006) ‘The structure-specific endonuclease Mus81-Eme1 promotes conversion of interstrand DNA crosslinks into double-strands breaks’, EMBO J, 25(20), pp. 4921–32.

Huang, Y., Pettitt, S. J., Guo, G., Liu, G., Li, M. A., Yang, F. and Bradley, A. (2012) ‘Isolation of homozygous mutant mouse embryonic stem cells using a dual selection system’, Nucleic Acids Res, 40(3), pp. e21.

Ip, S. C., Rass, U., Blanco, M. G., Flynn, H. R., Skehel, J. M. and West, S. C. (2008) ‘Identification of Holliday junction resolvases from humans and yeast’, Nature, 456(7220), pp. 357–61.

Kaemena, D. F., Yoshihara, M., Beniazza, M., Ashmore, J., Zhao, S., Bertenstam, M., Olariu, V., Katayama, S., Okita, K., Tomlinson, S. R., Yusa, K. and Kaji, K. (2023) ‘B1 SINE-binding ZFP266 impedes mouse iPSC generation through suppression of chromatin opening mediated by reprogramming factors’, Nat Commun, 14(1), pp. 488.

Leahy, J. J., Golding, B. T., Griffin, R. J., Hardcastle, I. R., Richardson, C., Rigoreau, L. and Smith, G. C. (2004) ‘Identification of a highly potent and selective DNA-dependent protein kinase (DNA-PK) inhibitor (NU7441) by screening of chromenone libraries’, Bioorg Med Chem Lett, 14(24), pp. 6083–7.

Lefort, N., Feyeux, M., Bas, C., Féraud, O., Bennaceur-Griscelli, A., Tachdjian, G., Peschanski, M. and Perrier, A. L. (2008) ‘Human embryonic stem cells reveal recurrent genomic instability at 20q11.21’, Nat Biotechnol, 26(12), pp. 1364–6.

Lilley, D. M. J. (2017) ‘Holliday junction-resolving enzymes-structures and mechanisms’, FEBS Lett, 591(8), pp. 1073–1082.

Livak, K. J. and Schmittgen, T. D. (2001) ‘Analysis of relative gene expression data using real-time quantitative PCR and the 2(-Delta Delta C(T)) Method’, Methods, 25(4), pp. 402–8.

Love, M. I., Huber, W. and Anders, S. (2014) ‘Moderated estimation of fold change and dispersion for RNA-seq data with DESeq2’, Genome Biol, 15(12), pp. 550.

Luo, G., Yao, M. S., Bender, C. F., Mills, M., Bladl, A. R., Bradley, A. and Petrini, J. H. (1999) ‘Disruption of mRad50 causes embryonic stem cell lethality, abnormal embryonic development, and sensitivity to ionizing radiation’, Proc Natl Acad Sci U S A, 96(13), pp. 7376–81.

Mayshar, Y., Ben-David, U., Lavon, N., Biancotti, J. C., Yakir, B., Clark, A. T., Plath, K., Lowry, W. E. and Benvenisty, N. (2010) ‘Identification and classification of chromosomal aberrations in human induced pluripotent stem cells’, Cell Stem Cell, 7(4), pp. 521–31.

Rantakari, P., Nikkilä, J., Jokela, H., Ola, R., Pylkäs, K., Lagerbohm, H., Sainio, K., Poutanen, M. and Winqvist, R. (2010) ‘Inactivation of Palb2 gene leads to mesoderm differentiation defect and early embryonic lethality in mice’, Hum Mol Genet, 19(15), pp. 3021–9.

Rass, U., Compton, S. A., Matos, J., Singleton, M. R., Ip, S. C., Blanco, M. G., Griffith, J. D. and West, S. C. (2010) ‘Mechanism of Holliday junction resolution by the human GEN1 protein’, Genes Dev, 24(14), pp. 1559–69.

Savatier, P., Lapillonne, H., Jirmanova, L., Vitelli, L. and Samarut, J. (2002) ‘Analysis of the cell cycle in mouse embryonic stem cells’, Methods Mol Biol, 185, pp. 27–33.

Spits, C., Mateizel, I., Geens, M., Mertzanidou, A., Staessen, C., Vandeskelde, Y., Van der Elst, J., Liebaers, I. and Sermon, K. (2008) ‘Recurrent chromosomal abnormalities in human embryonic stem cells’, Nat Biotechnol, 26(12), pp. 1361–3.

Stead, E., White, J., Faast, R., Conn, S., Goldstone, S., Rathjen, J., Dhingra, U., Rathjen, P., Walker, D. and Dalton, S. (2002) ‘Pluripotent cell division cycles are driven by ectopic Cdk2, cyclin A/E and E2F activities’, Oncogene, 21(54), pp. 8320–33.

Tang, Z., Liang, Z., Zhang, B., Xu, X., Li, P., Li, L., Lu, L. Y. and Liu, Y. (2024) ‘MRE11 is essential for the long-term viability of undifferentiated spermatogonia’, Cell Prolif, 57(9), pp. e13685.

Tichy, E. D., Pillai, R., Deng, L., Liang, L., Tischfield, J., Schwemberger, S. J., Babcock, G. F. and Stambrook, P. J. (2010) ‘Mouse embryonic stem cells, but not somatic cells, predominantly use homologous recombination to repair double-strand DNA breaks’, Stem Cells Dev, 19(11), pp. 1699–711.

Tsuzuki, T., Fujii, Y., Sakumi, K., Tominaga, Y., Nakao, K., Sekiguchi, M., Matsushiro, A., Yoshimura, Y. and Morita T (1996) ‘Targeted disruption of the Rad51 gene leads to lethality in embryonic mice’, Proc Natl Acad Sci U S A, 93(13), pp. 6236–40.

Wang, X., Wang, H., Guo, B., Zhang, Y., Gong, Y., Zhang, C., Xu, H. and Wu, X. (2016) ‘Gen1 and Eme1 Play Redundant Roles in DNA Repair and Meiotic Recombination in Mice’, DNA Cell Biol, 35(10), pp. 585–590.

Wechsler, T., Newman, S. and West, S. C. (2011) ‘Aberrant chromosome morphology in human cells defective for Holliday junction resolution’, Nature, 471(7340), pp. 642–6.

White, J., Stead, E., Faast, R., Conn, S., Cartwright, P. and Dalton, S. (2005) ‘Developmental activation of the Rb-E2F pathway and establishment of cell cycle-regulated cyclin-dependent kinase activity during embryonic stem cell differentiation’, Mol Biol Cell, 16(4), pp. 2018–27.

Wu, L. and Hickson, I. D. (2003) ‘The Bloom’s syndrome helicase suppresses crossing over during homologous recombination’, Nature, 426(6968), pp. 870–4.

Wyatt, H. D. and West, S. C. (2014) ‘Holliday junction resolvases’, Cold Spring Harb Perspect Biol, 6(9), pp. a023192.

Xiao, Y. and Weaver, D. T. (1997) ‘Conditional gene targeted deletion by Cre recombinase demonstrates the requirement for the double-strand break repair Mre11 protein in murine embryonic stem cells’, Nucleic Acids Res, 25(15), pp. 2985–91.

Yoon, S. W., Kim, D. K., Kim, K. P. and Park, K. S. (2014) ‘Rad51 regulates cell cycle progression by preserving G2/M transition in mouse embryonic stem cells’, Stem Cells Dev, 23(22), pp. 2700–11.

Zhu, J., Petersen, S., Tessarollo, L. and Nussenzweig, A. (2001) ‘Targeted disruption of the Nijmegen breakage syndrome gene NBS1 leads to early embryonic lethality in mice’, Curr Biol, 11(2), pp. 105–9.

