## Supplemental figures for "The Holliday junction resolvase GEN1 preserves genome integrity and self-renewal in mouse embryonic stem cells"

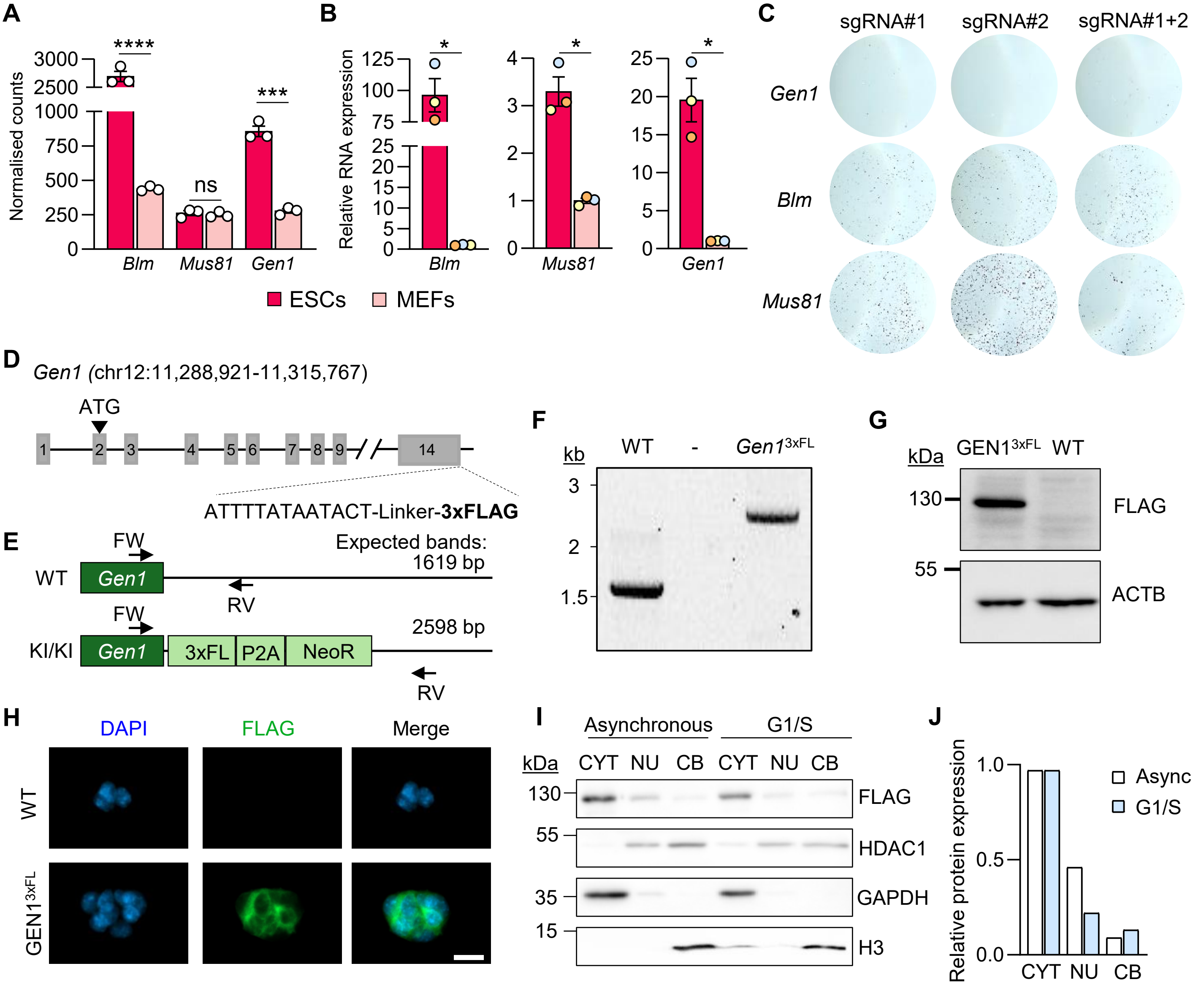


**Figure S1.** (A) RNA Expression of *Mus81, Gen1* and *Blm* in ESCs versus mouse embryonic stem cells (MEFs) using normalised counts from a published RNA-seq dataset (GSE198166) and (B) in cDNA samples from ESCs relative to Gapdh and MEF samples (n=3 biological replicates). (C) Representative images of wells after AP staining at day 3 post-transfection of CRISPR/Cas9 construct to generate *Gen1*, *Blm* and *Mus81* KOs. (D) Schematic of the *Gen1* genomic locus showing the insertion site of the 3×FLAG tag by knock-in (KI). (E) Schematic of the genotyping PCR strategy used to distinguish KI/KI from wild-type (WT) ESCs. (F) Representative PCR products from wild-type and GEN1$^{3\times FL}$ knock-in (KI) ESCs. (G) Western blot of whole-cell extracts from GEN1$^{3\times FL}$ ESCs. Antibodies used: anti-FLAG (GEN1) and anti-β-actin (ACTB; loading control). (H) Immunofluorescence staining with anti-FLAG antibody in WT and GEN1$^{3\times FL}$ ESCs. Nuclei were counterstained with DAPI. Scale bar: 20 µm. (I) Western blot of subcellular fractions from asynchronous and G1/S-synchronized GEN1$^{3\times FL}$ ESCs. Antibodies against FLAG (tag for GEN1), HDAC1 (marker for nucleus, NU), GAPDH (marker for cytoplasm, CYT) and H3 (marker for chromatin-bound fraction, CB) were used. (J) Quantification of the protein levels from (I) of GEN1 at the different compartments relative to asynchronous samples. Statistical significance was assessed by unpaired t-test relative to MEFs conditions. * $p < 0.05$; *** $p < 0.001$; **** $p < 0.0001$; ns: non-significant.

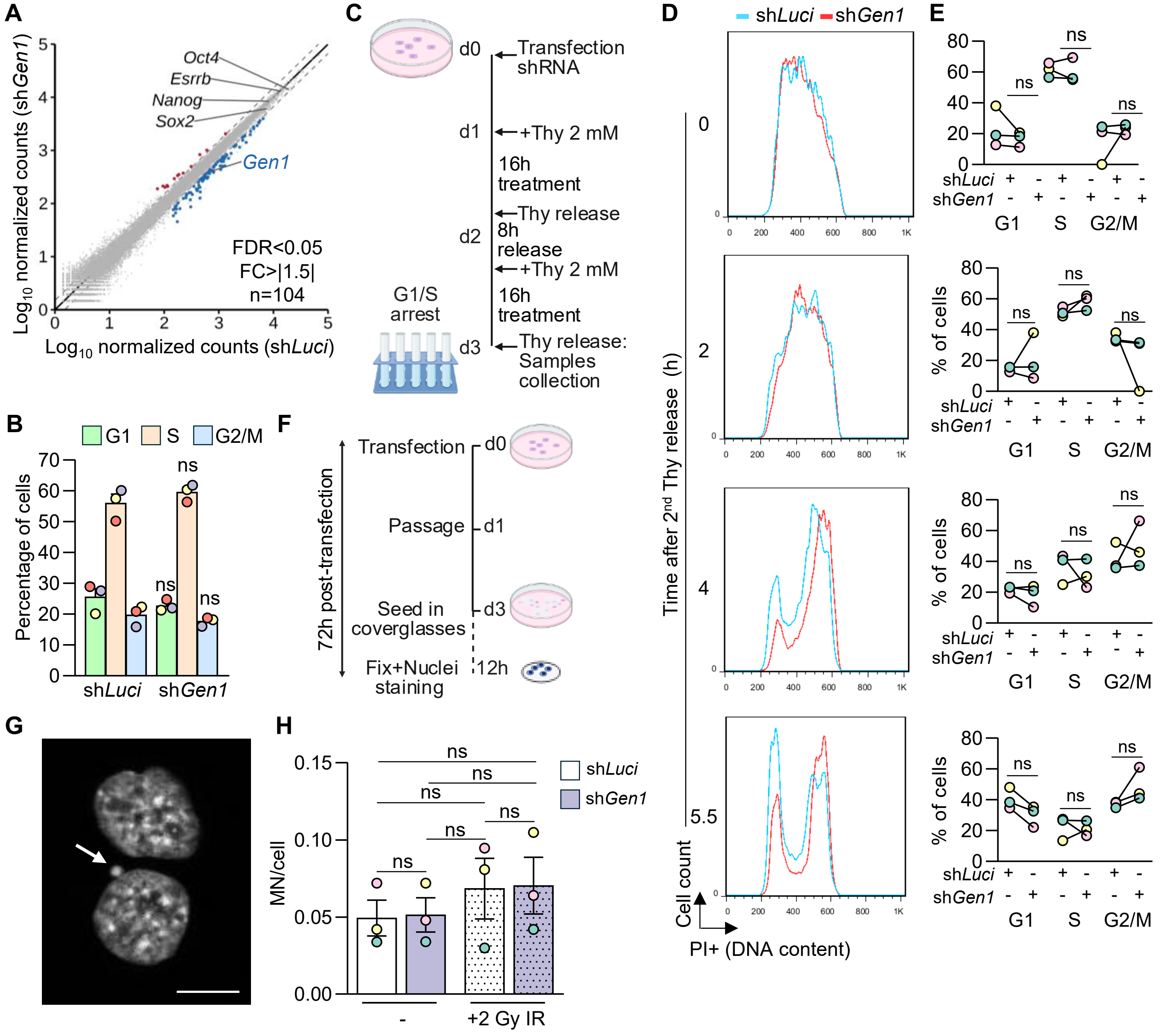


**Figure S2.** (A) Scatter plot of gene expression values (expressed as $Log_{10}$ normalised counts) obtained by RNA-seq analysis comparing the expression of sh*Luci* vs sh*Gen1*. Dots corresponding to expression levels of *Gen1* and *Oct4*, *Esrrb*, *Nanog* and *Sox2* pluripotency markers are indicated. (B) Quantification of the percentage of cells in G1, S, and G2/M phases after flow cytometry analysis. Data represent mean ± SEM of n = 3 biological replicates. (C) Scheme for the experimental approach followed for the synchronization of cells in G1/S with a double thymidine (Thy) block after transfection of shRNAs against *Gen1 (*sh*Gen1)* and *Luci* as control. Samples were collected at different time points after the second Thy release. (D) Representative histograms showing cell cycle progression of ESCs through the different stages. The blue line corresponds to sh*Luci* population and the red line to sh*Gen1* population. (E) Quantification of the percentage of cells in G1, S, and G2/M phases under *Luci* and *Gen1* knockdown conditions (n = 3, biological replicates, each colour represents an independent experiment) at 0 h, 2 h, 4 h and 5.5 h after the second Thy release. Statistical significance was determined by unpaired t-test: ns: non-significant. (F) Schematic of the experimental approach for detection of micronuclei formation. (G) Representative images of a cell without (upper) and with (lower) micronucleus (white arrow). Scale bar: 10 µm. (H) Quantification of micronuclei (MN) per cell at 72 h post-transfection with shRNAs, with or without 2 Gy ionizing radiation (IR) treatment (n = 3, biological replicates). Statistical significance was assessed by unpaired two-tailed t-test. ns, non-significant.

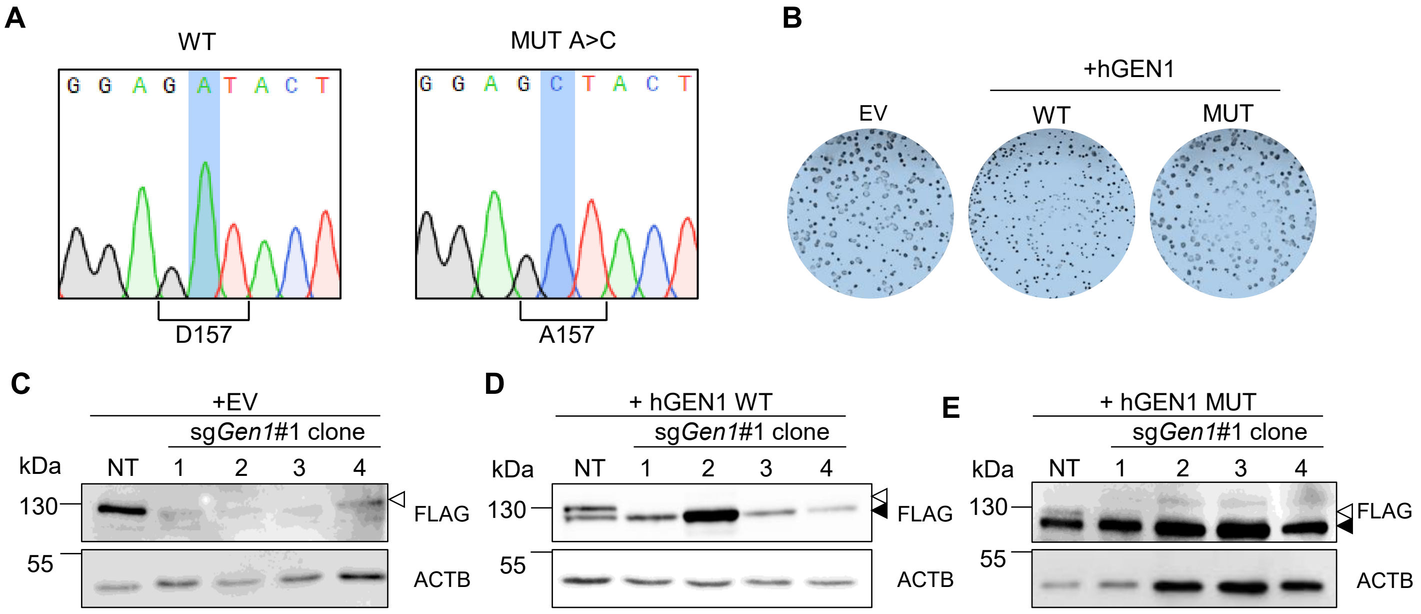


**Figure S3.** (A) Sanger sequencing confirming the point mutation generating the catalytically inactive human GEN1 mutant (D157A). (B) Representative images of CFA wells from ESCs with or without overexpression of hGEN1 WT or MUT or EV control. Western blots of selected clones following *Gen1* KO in GEN1$^{3\times FL}$ + EV ESCs (C), GEN1$^{3\times FL}$ + hGEN1 WT ESCs (D), and GEN1$^{3\times FL}$ + hGEN1 MUT (E) ESCs. Anti-FLAG antibody detects both endogenous mouse GEN1 (upper band, white arrowhead) and human GEN1 (lower band, black arrowhead). ACTB was used as loading control.

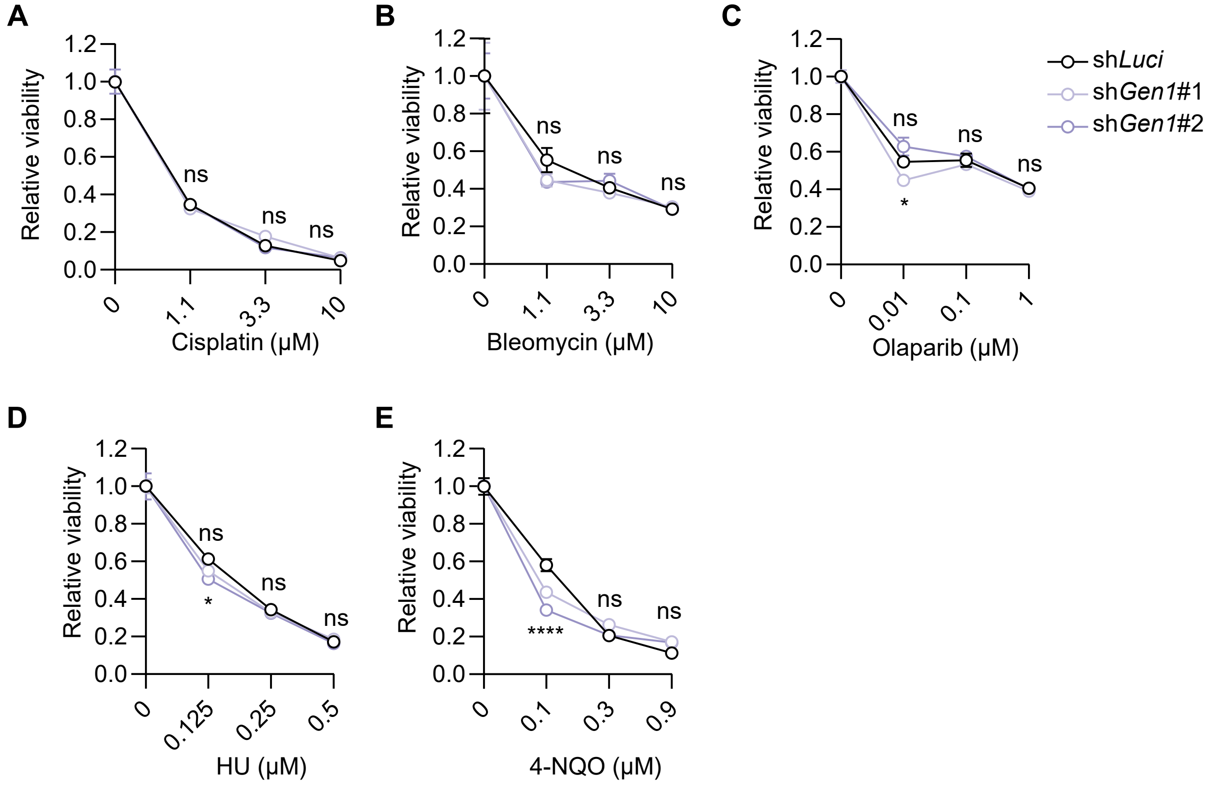


**Figure S4.** Relative viability (normalized to samples treated with vehicle at 0 μM) of sh*Luci* and sh*Gen1* ESCs (two independent shRNAs) treated with increasing concentrations of (A) cisplatin, (B) bleomycin, (C) olaparib, (D) hydroxyurea (HU) and (E) 4-nitroquinoline-1-oxide (4-NQO). Statistical significance was assessed by two-way ANOVA with Dunnett's multiple comparisons test, using sh*Luci* as the reference group. *$p < 0.05$; ****$p < 0.0001$; ns, non-significant.
